# Phenotypic screening of covalent compound libraries identifies chloromethyl ketone antibiotics and MiaA as a new target

**DOI:** 10.1101/2024.01.22.576576

**Authors:** Yizhen Jin, Sadhan Jana, Mikail E. Abbasov, Hening Lin

## Abstract

The emerging antibiotic resistance requires the development of new antibiotics working on novel bacterial targets. Here, we reported an antibiotic discovery workflow by combining the cysteine-reactive compound library phenotypic screening with activity-based protein profiling, which enables the rapid identification of lead compounds as well as new druggable targets in pathogens. Compounds featuring chloromethyl ketone scaffolds exhibited a notably high hit rate against both gram-negative and gram-positive bacterial strains, but not the more commonly used warheads such as acrylamide or chloroacetamide. Target identification of the lead compound, 10-F05, revealed that its primary targets in *S. flexneri* are FabH Cys112 and MiaA Cys273. We validated the target relevance through biochemical and genetic interactions. Mechanistic studies revealed modification of MiaA by 10-F05 impair substrate tRNA binding, leading to decreased bacterial stress resistance and virulence. Our findings underscore chloromethyl ketone as a novel antibacterial warhead in covalent antibiotic design. The study showcases that combining covalent compound library phenotypic screening with chemoproteomics is an efficient way to identify new drug targets as well as lead compounds, with the potential to open new research directions in drug discovery and chemical biology.

**Graphic Abstract:** 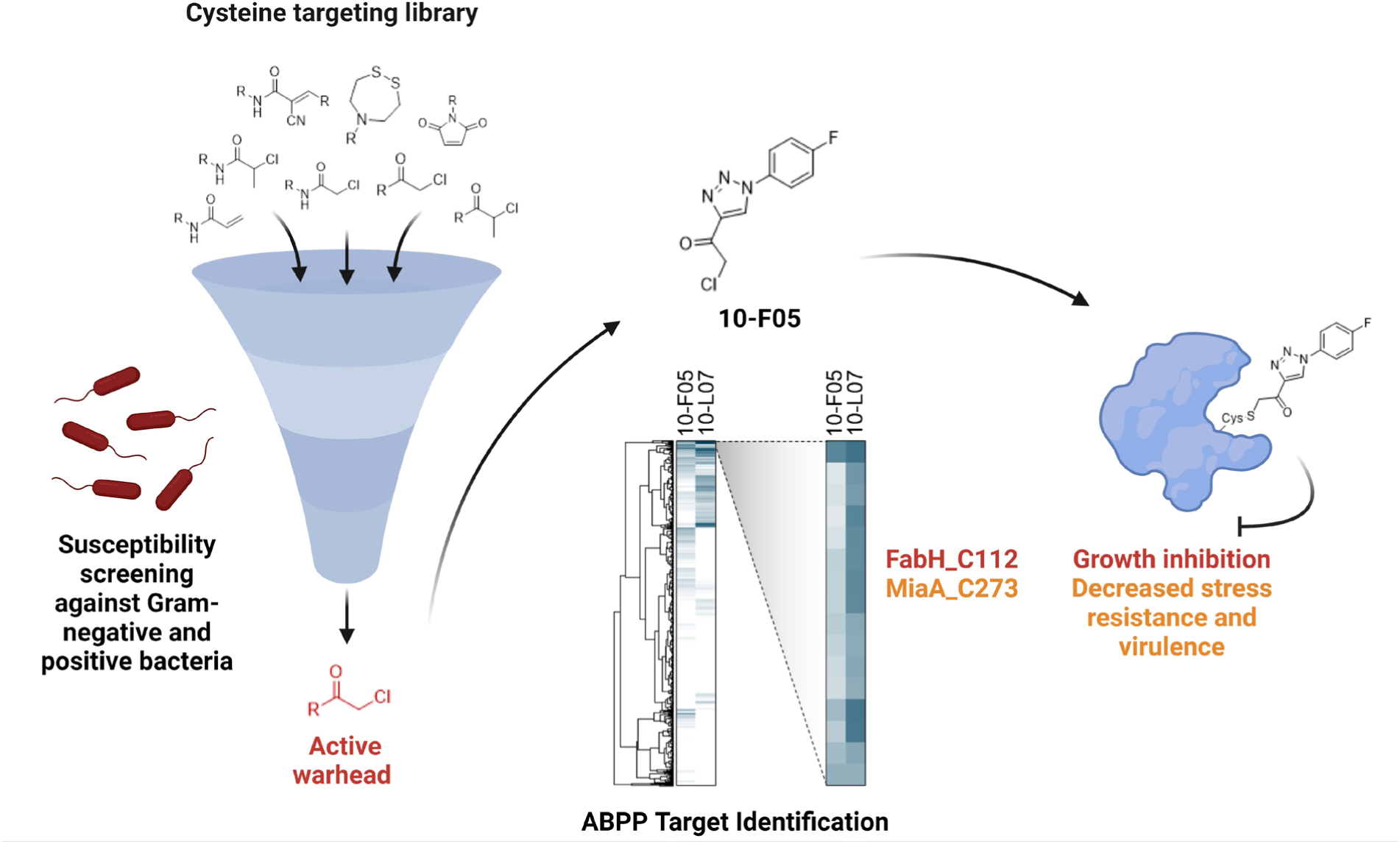

## Introduction

Antibiotics are essential human medicines, but antibiotics resistance is threatening the effectiveness of existing antibiotics. Expanding the targets pool is crucial to overcome antibiotics-resistance. Known antibiotics target a limited number of well-studied targets, including the biosynthesis of cell wall, proteins, and nucleic acids.^1^ There is a need to explore novel strategies that can target a broader range of bacterial proteins. Inspired by the recent successful development of KRAS G12C inhibitor to target proteins that traditionally considered as ‘undruggable’^2^ and the existence of antibiotics that covalently target proteins (such as penicillin), we hypothesized that screening a cysteine-reactive compound library can expand the pool of druggable proteins in bacterial proteomes. By utilizing covalent interactions, this approach has the potential to engage essential proteins that are untouched in conventional drug screenings. Moreover, the use of chemical proteomics, taking advantage of the covalent nature of the hit compounds, can significantly streamline the challenging process of target elucidation.^3–5^ By combining these methodologies, we can accelerate the identification and characterization of novel targets, paving the way for the development of new antibiotics.

Despite the rising interest in cysteine targeting compounds,^6–8^ previous studies primarily used very small libraries with a limited number of cysteine warheads like chloroacetamide.^9,10^ The limitation of warhead diversity could hinder the identification of druggable protein targets. The diverse properties and reactivities of bacterial cysteinome may require a broader range of warheads to effectively interact with and modify target proteins. Here, we applied a larger and more diverse cysteine-targeting library (Cys-library) consisting of 3,200 fragment-like covalent ligands in antibacterial screening to discover potential new antibiotics and new targets. We discovered that chloromethyl ketone scaffolds showed promising activity in both gram-positive and gram-negative strains. One lead compound, 10-F05, displays broad-spectrum antibacterial activity. Using competitive activity-based protein profiling (ABPP), we identified and validated two proteins, FabH and MiaA, as the physiologically relevant targets of 10-F05. Importantly, MiaA is not a previously recognized antibiotics target, underscoring the advantage of employing phenotypic screening of covalent compound libraries with ABPP to identify both lead compounds and novel targets.

## Results

### Susceptibility screenings identify antibacterial hits with chloromethyl ketone warhead

To expand the druggable targets within bacterial cysteinome, we obtained a diverse Cys-library consisting of 3,200 fragment-like compounds from Enamine (**Figure S1A-E**). We first assessed the cytotoxicity of the Cys-library in HEK293T cells at a concentration of 25 μM over two days (**Figure S2**). Chloromethyl ketone and chloroacetamide scaffolds exhibited higher cytotoxicity in HEK293T cells compared to other warhead scaffolds. The 2-chloropropinoamide and 2-chloroethyl ketone scaffolds displayed higher tolerance in the mammalian cells.

Next, we screened the Cys-library to identify active compounds against *S. aureus* (MSSA476) and *V. cholerae* (SAD30) at 25 μM (**Figure 1A** and **1B**). From the initial screening, we identified 48 and 17 hits that inhibited *S. aureus* and *V. cholerae* growth in liquid culture, respectively (**Table S1**). Interestingly, many *S. aureus* active hits were not active in *V. cholerae*, which could be due to differences of cell wall composition or cysteine targets in these two strains.^10,11^ We excluded 11 chloroacetamides, three α-cyanoacrylamides, and a cluster of dimethyl pyrroles (DP) with chloromethyl ketone warhead due to the high cytotoxicity in HEK293T cells at 25 μM (**Figure S3B, S3C, S3D,** and **supplementary table**).^10,12^ Two compounds (3-J04 and 8-I07, **Figure S3A**) were excluded from further analysis due to the presence of a nitrofuran moiety, which is known to be toxic.^13^ In total,15 chloromethyl ketone compounds were found to be active against both *S. aureus* and *V. cholerae*. These hits and three additional chloromethyl ketones were selected to measure the minimum inhibitory concentrations (MIC) against an expanded collection of pathogens including *S. flexneri* M90T, *V. cholerae* SAD30, *E. coli* JPN15, and *S. aureus* MSSA476 (**Table 1**). Most chloromethyl ketone hits displayed MICs ranging from 5 μM to 50 μM, with slight variations observed among the four pathogens.

**Figure 1.**
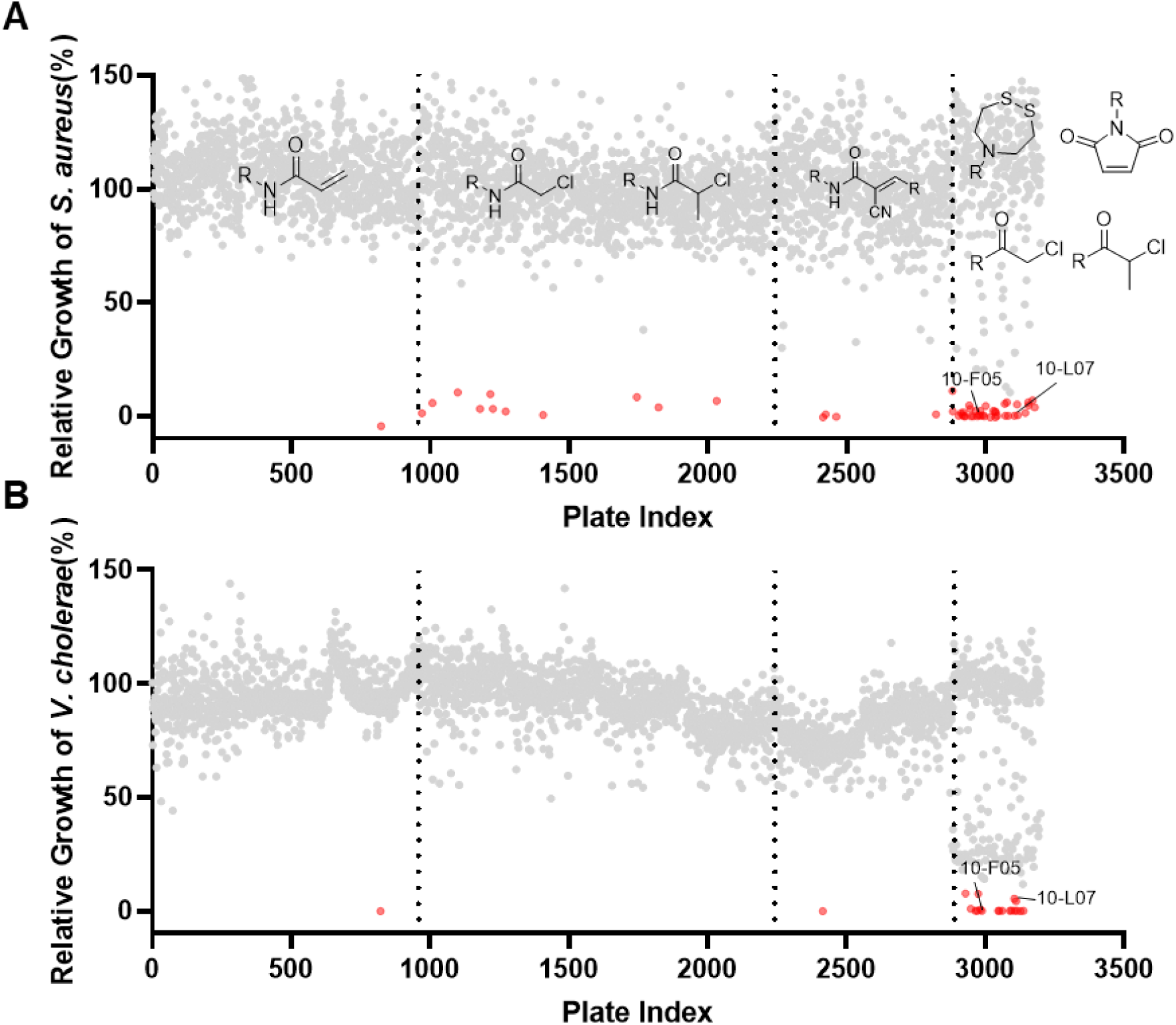
Bacterial growth inhibition screenings of a Cys-library at 25 µM. Relative bacterial growth was determined by normalizing the OD600 value to the DMSO control. (**A**) Screenings using *S. aureus* (MSSA476). (**B**) Screenings using *V. cholerae* (SAD30). Hits (relative growth ≤ 10%) are labeled in red.

**Table 1.**
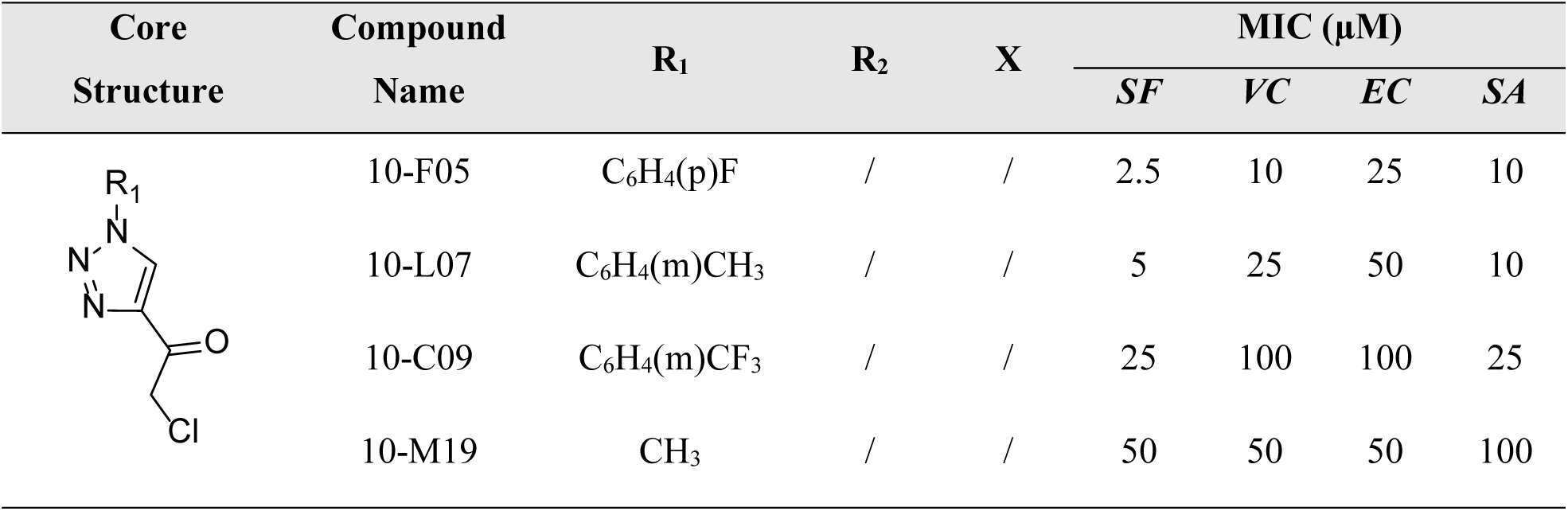
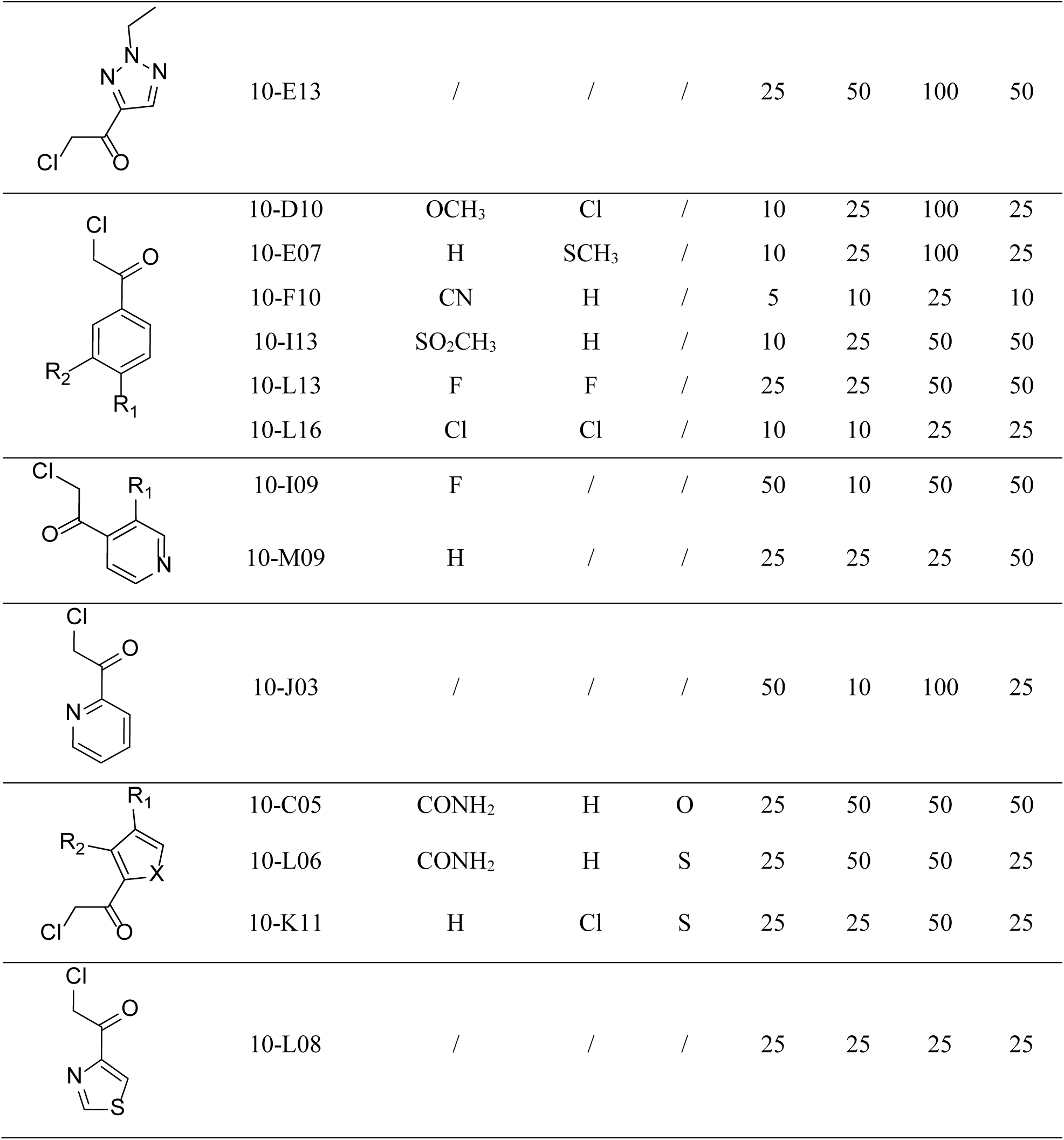
MIC values of hits with chloromethyl ketone warhead.

| Core Structure | Compound Name | R<sub>1</sub> | R<sub>2</sub> | X | MIC (μM) | | | |
| --- | --- | --- | --- | --- | --- | --- | --- | --- |
|  |  |  |  |  | SF | VC | EC | SA |
|  | 10-F05 | C<sub>6</sub>H<sub>4</sub>(p)F | / | / | 2.5 | 10 | 25 | 10 |
|  | 10-L07 | C<sub>6</sub>H<sub>4</sub>(m)CH<sub>3</sub> | / | / | 5 | 25 | 50 | 10 |
|  | 10-C09 | C<sub>6</sub>H<sub>4</sub>(m)CF<sub>3</sub> | / | / | 25 | 100 | 100 | 25 |
|  | 10-M19 | CH<sub>3</sub> | / | / | 50 | 50 | 50 | 100 |
|  | 10-E13 | / | / | / | 25 | 50 | 100 | 50 |
|  | 10-D10 | OCH<sub>3</sub> | Cl | / | 10 | 25 | 100 | 25 |
|  | 10-E07 | H | SCH<sub>3</sub> | / | 10 | 25 | 100 | 25 |
|  | 10-F10 | CN | H | / | 5 | 10 | 25 | 10 |
|  | 10-I13 | SO<sub>2</sub>CH<sub>3</sub> | H | / | 10 | 25 | 50 | 50 |
|  | 10-L13 | F | F | / | 25 | 25 | 50 | 50 |
|  | 10-L16 | Cl | Cl | / | 10 | 10 | 25 | 25 |
|  | 10-I09 | F | / | / | 50 | 10 | 50 | 50 |
|  | 10-M09 | H | / | / | 25 | 25 | 25 | 50 |
|  | 10-J03 | / | / | / | 50 | 10 | 100 | 25 |
|  | 10-C05 | CONH<sub>2</sub> | H | O | 25 | 50 | 50 | 50 |
|  | 10-L06 | CONH<sub>2</sub> | H | S | 25 | 50 | 50 | 25 |
|  | 10-K11 | H | Cl | S | 25 | 25 | 50 | 25 |
|  | 10-L08 | / | / | / | 25 | 25 | 25 | 25 |

### Appropriate thiol reactivity is required for high antibacterial activity

Surprisingly, the two most frequently used cysteine covalent warheads,^14^ acrylamide and chloroacetamide, were not found among *V. cholerae* active hits despite that they account for the majority of the Cys-library (**Table S1**). Only 11 out of 751 chloroacetamide compounds were active against *S. aureus* and none against *V. cholerae*. A previous study also showed that several chloroacetamide-based MurA covalent inhibitors (47 fragment-sized chloroacetamide scaffolds) were not active against *E. coli*.^9^ In contrast, a less studied warhead, chloromethyl ketone, has the highest hit discovery rate in both *V. cholerae* and *S. aureus* screenings.

Chloromethyl ketone is not commonly employed in cysteine-targeting compound libraries, but there are precedents in enzyme inhibitors. One example is tosyl phenylalanyl chloromethyl ketone (TPCK), which covalently binds to the active site cysteine of Caspase-3.^15^ The warhead is crucial for forming the covalent bond with the target protein. The reaction rate constant (*kinact*) is mainly contributed by the intrinsic reactivity of warhead.^6^ However, there had been no systematic studies comparing the reactivities of the covalent warheads used in this study. Thus, to profile the reactivity difference, we employed a reduced 5,5′ -dithiobis-(2-nitrobenzoic acid) (DTNB, Ellman’s reagent) assay (**Figure S4A** and **Figure 2A**).^16^ Most compounds could be fitted into a second-order reaction model (**Figure S4B**). Chloromethyl ketone exhibited the highest reactivity among all the warheads tested, with a *k*avg of 4.59 × 10^-6^ M^−1^ s^−1^, which was approximately 1.2-fold higher than maleimide (*k*avg = 3.59 × 10^−6^ M^−1^ s^−1^), 6.2-fold higher than chloroacetamide (*k*avg = 7.45 × 10^−7^ M^−1^ s^−1^), 17-fold higher than acrylamide (*k*avg = 2.69× 10^−7^ M^−1^ s^−1^). Chloroethyl ketone, which has an extra methyl group at α position, displayed decreased reactivity (*k*avg = 1.88 × 10^-6^ M^-1^ s^-1^). Disulfide and α-cyanoacrylamide are not tested in this assay due to the TCEP (tris(2-carboxyethyl) phosphine) interference of disulfide bond and reversible covalent nature of α-cyanoacrylamide.

**Figure 2.**
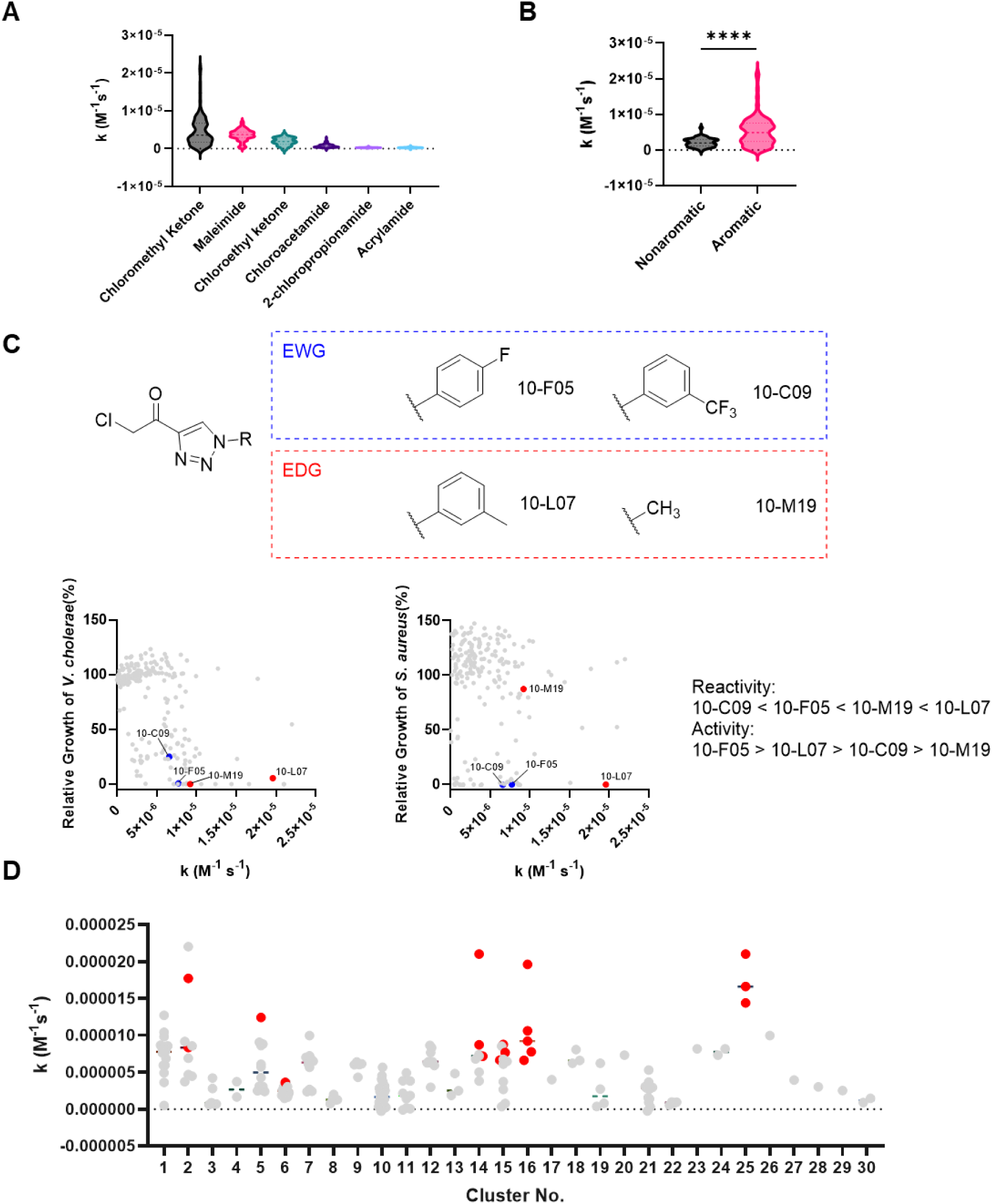
Suitable reactivity of warhead is important for antibacterial activity. (**A**) Thiol reactivity of compounds containing different warheads in the Cys-library measured using reduced DTNB. (**B**) Comparison of thiol reactivity of chloromethyl ketones with and without aromatic ring conjugated. (**C**) Top, structure of 10-F05 and its derivatives. Bottom, the reactivity of 210 chloromethyl ketone compounds is plotted against the relative growth of *V. cholerae* and *S. aureus*. (**D**) Comparison of thiol reactivity in different chloromethyl ketone clusters. Compounds are clustered based on structural similarity in Data Warrior. Red ones are identified as antibacterial hits in Table 1. The p values (unpaired two-tailed t-tests) were calculated in GraphPad Prism (version 9.3.1). ****p < 0.0001.

Notably, all active hits possessed a direct conjugation of the chloromethyl ketone warhead with an aromatic ring. The aromatic ring conjugated chloromethyl ketone compounds are generally more reactive than the nonaromatic ones because of the conjugation effect (**Figure 2B**). Both 10-C09 and 10-F05 possess electron-withdrawing groups (trifluoromethyl and fluoro, respectively, **Figure 2C**) on the benzene ring, which weaken the conjugation effect, leading to decreased reactivity compared to 10-L07, which has an electron-donating group on the benzene ring (**Figure 2C)**. Interestingly, the thiol reactivity did not correlate with the antibacterial activity. For instance, 10-C09 and 10-F05 exhibited similar reactivity towards thiols, yet they displayed a 2.5∼10-fold difference in their MICs across four tested pathogens, as shown in **Table 1**. Furthermore, the two derivatives of 10-F05, namely 10-M19 and 10-L07, which are 1.2-fold and 2.5-fold more reactive, respectively, exhibited 2∼10-fold worse MICs compared to 10-F05. These results emphasize that other factors contribute to the MIC values.

To better understand the structure activity relationship (SAR), we employed t-SNE distribution in Data Warrior to cluster the 210 compounds in the Cys-library that contained a chloromethyl ketone warhead (**Figure 2D** and **Figure S5**).^17^ Totally 30 clusters were assigned based on the structural similarity. Despite the important role of high reactivity plays in antibacterial activity, no hits were observed in several highly reactive structure clusters 1, 7, and 12, emphasizing the significance of non-covalent moieties in facilitating target engagement. We propose that the high reactivity of 10-F05 is necessary to achieve rapid target engagement, but reactivity is not the sole determining factor for its antibacterial activity.

### 10-F05 is active against multiple ESKAPE pathogens and has slow resistance development

We carried further experiments with 10-F05 and found that 10-F05 inhibits multiple ESKAPE pathogens, including methicillin resistant *S. aureus* (MRSA), *E. cloacae*, *K. pneumoniae*, and *E. coli*. 10-F05 lost its activity in several multiple drug resistant Gram-negative strains (**Table 2**, antibiotic spectrum for bacterial strains used is reported in **Supplementary Data**). A time-killing experiment was conducted using a starting inoculum of approximately 10^5^ CFU/mL in LB medium. The results demonstrated that 2.5 µM of 10-F05 was able to eliminate ∼99% of *S. flexneri* within a 24-hour period (**Figure 3A**). Furthermore, 10-F05 exhibited low cytotoxicity in HEK293T cells when compared to other chloromethyl ketone hits (**Figure S6A**). The IC50 value of 10-F05 is ∼50 µM in both HEK293T and A549 cells after 2 days incubation. The selectivity ratio ranged from 2 to 20 in different strains (**Figure S6B**).

**Figure 3.**
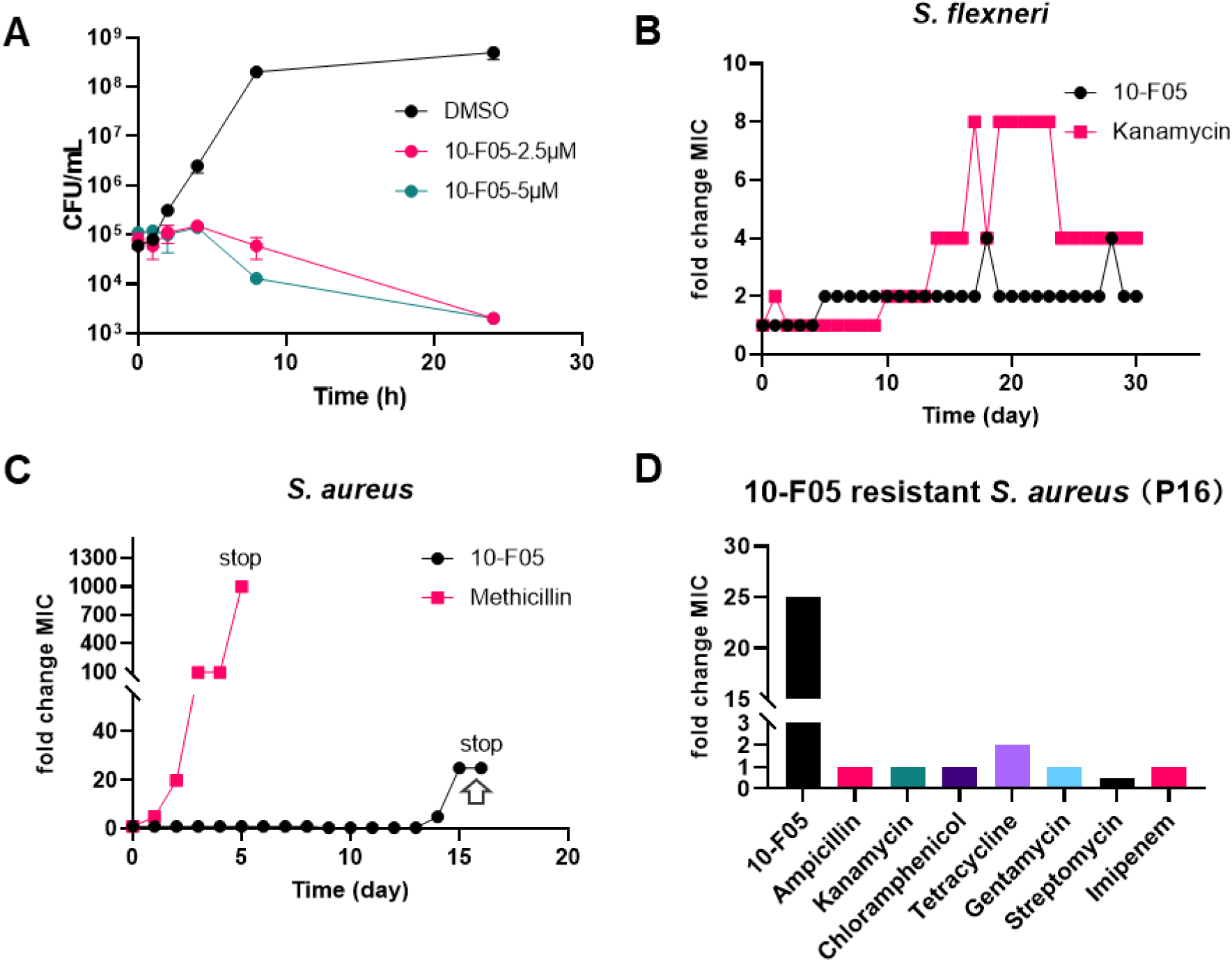
Antibacterial activity and resistance development of 10-F05. (**A**) Time-dependent killing of *S. flexneri* M90T (N=2). (**B**) Resistance development of *S. flexneri* M90T against 10-F05 and Kanamycin during daily serial passaging with sub-MIC concentrations. (C) Resistance development of *S. aureus* MSSA476 against 10-F05 and Methicillin during daily serial passaging with sub-MIC concentrations. (**D**) MIC fold change of 10-F05 resistant *S. aureus* MSSA476 against other classes of antibiotics. MIC values were reported in Supplementary table.

**Table 2.**
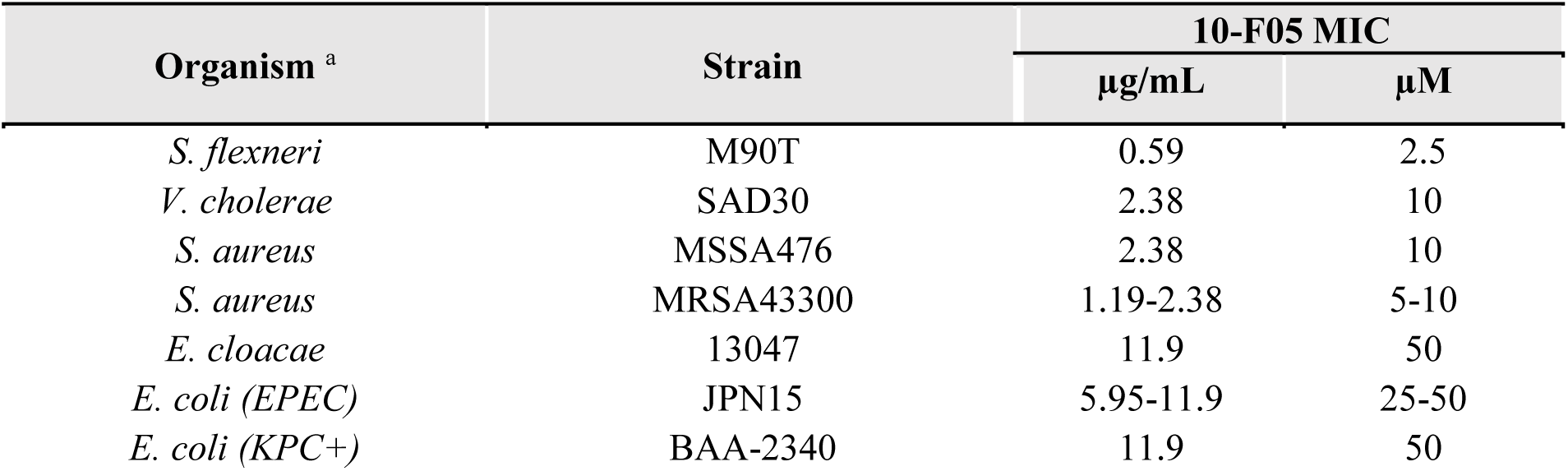
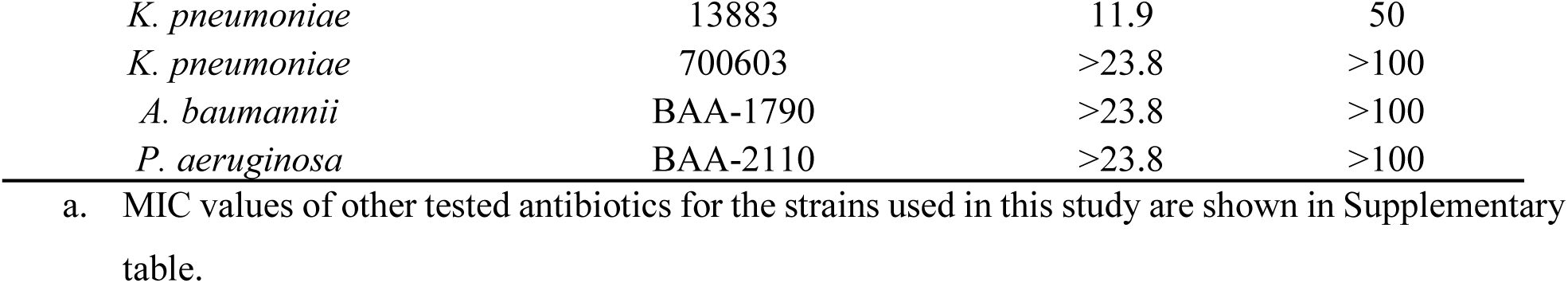
MIC of 10-F05 against an expanded bacterial panel.

Next, we tested the development of resistance against 10-F05 in *S. aureus* MSSA476 and *S. flexneri* M90T (**Figure 3B** and **3C**). *S. flexneri* and *S. aureus* were passaged daily with sub-MIC concentrations of 10-F05 or two known antibiotics (kanamycin and methicillin), and the surviving bacteria in the next day were used to measure the MIC values of the compounds. *S. flexneri* M90T exhibited a 2-fold increase in MIC to 10-F05 on day 5 (MIC = 5 µM, 2-fold change) and maintained less than 4-fold change of MIC after 30 days of sub-MIC daily passage. During this period, *S. flexneri* M90T developed higher resistance to kanamycin, with a maximum fold change in MIC of ≤ 8. In contrast, *S. aureus* gained resistance to methicillin with a MIC fold change higher than 200 after 5 days (MIC > 1 mg/mL). The resistance of *S. aureus* to 10-F05 occurred at a much slower rate, taking place on day 14 and finally reached a 25-fold change in MIC by day 16. The slow resistance development rate in both *S. flexneri* and *S. aureus* suggested a potential poly-pharmacological mechanism of 10-F05. Notably, the 10-F05 resistant *S. aureus* strains did not exhibit significant cross-resistance to other commonly used antibiotic classes, indicating a novel mechanism of action of 10-F05 (**Figure 3D**).

### Competitive ABPP identifies FabH, MiaA and PdxY as targets

Encouraged by its promising activity in both gram-positive and gram-negative bacteria, we selected 10-F05 as the lead compound for further studies. To confirm that 10-F05 inhibits bacterial growth by targeting the bacterial cysteinome, we measured the MICs of negative control compounds, in which the cysteine-reactive warheads were replaced with unreactive ones, against model pathogens (**Table S3**). All the negative control compounds showed no activity, suggesting that the growth inhibitory effect is through the covalent modification of bacterial proteins.

10-F05 and 10-L07 exhibited the best MICs of 2.5 and 5 µM, respectively, in *S. flexneri* M90T and they are quite similar in structure despite their 2.2-fold difference in thiol reactivity. We hypothesized that they engage similar cysteines to inhibit bacterial growth. We applied competitive activity-based protein profiling (ABPP) in *S. flexneri* M90T using both 10-F05 and 10-L07 to increase the target confidence. Live *S. flexneri* M90T cells suspended in PBS were treated with 10-F05, 10-L07, or DMSO and lysed, followed by treating with iodoacetamide-conjugated desthiobiotin (IA-DTB). After trypsin and Lys-C digestion, IA-DTB labeled peptides were affinity enriched and quantified using quantitative tandem mass tag (TMT) proteomics. Real-time search coupled with synchronous precursor selection and MS3 fragmentation were implemented to optimize both quantification and proteomic coverage (**Figure 4A**).^18^

**Figure 4.**
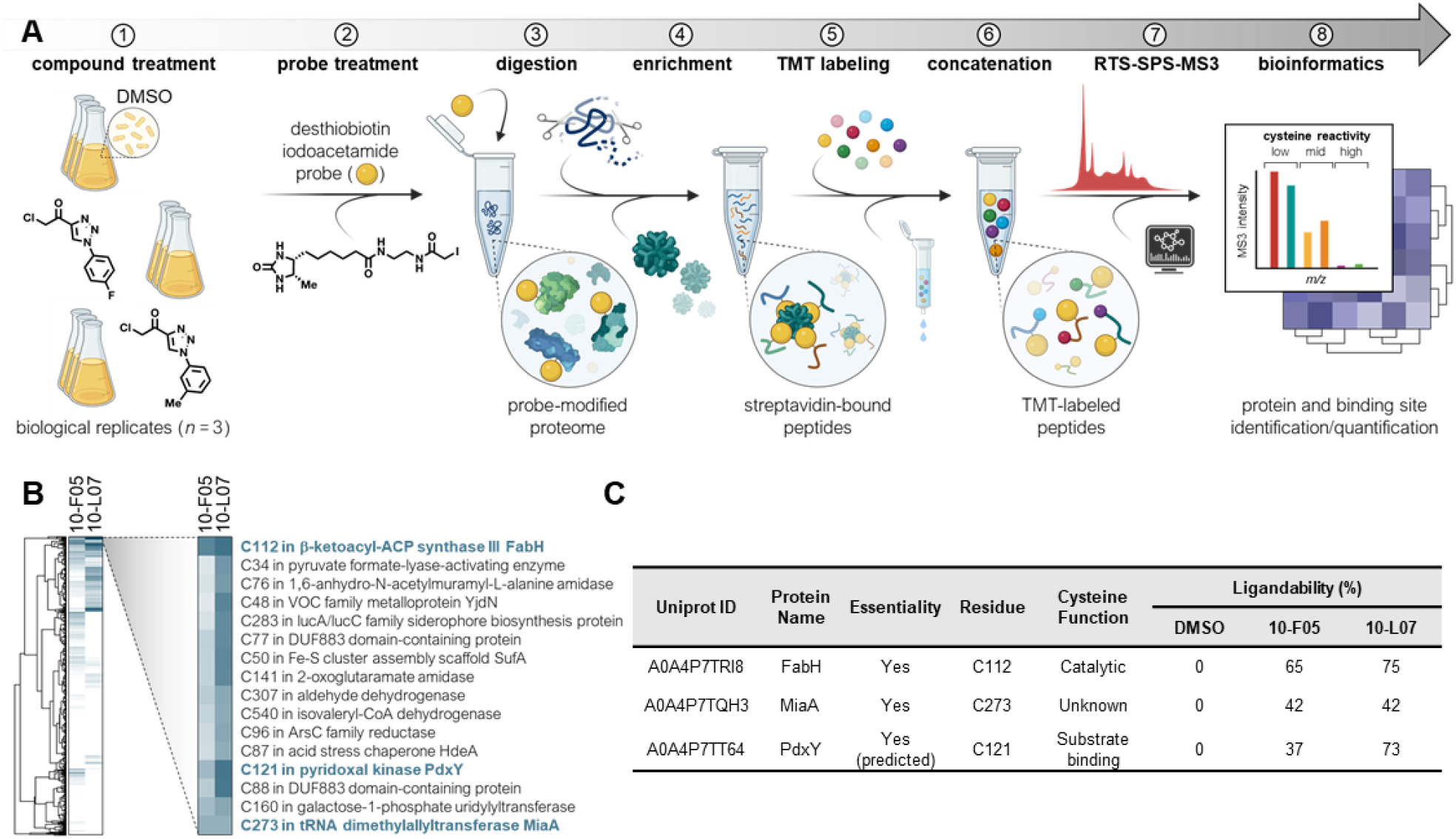
Competitive activity-based protein profiling identified FabH, MiaA and PdxY as targets of 10-F05. (**A**) Schematic of TMTpro-18plex-based workflow for mapping cysteine-drug interactions in live *S. flexneri*. (**B**) Heatmap analysis of the protein targets of 10-F05 and 10-L07. (**C**) Ligandability table of highlighted cysteines in **B**. Cysteine function is annotated based on Uniprot annotation.

In our proteomic result, we identified 1035 cysteines from 588 proteins in *S. flexneri* M90T. Gene ontology was assigned to the identified proteins and no obvious enrichment was observed (**Figure S7**). We next annotated all the identified proteins using NetGenes database to predict the essentiality of the identified proteins.^19^ Among 588 proteins, 103 were predicted to be essential in *S. flexneri* M90T (**Figure S8**).

We then quantified the cysteines that are engaged by the two compounds. As anticipated, 10-L07 engaged many cysteines with high ligandability due to its higher reactivity. However, 10-F05 was not as promiscuous as we expected and only engaged 5 cysteines higher than 35% ligandability in the bacterial proteome. Interestingly, FabH Cys112 was identified as the top cysteine hit in both 10-F05 and 10-L07, with a ligandability of 65% and 75%, respectively (**Figure 4B** and **4C**). FabH [β-ketoacyl-acyl carrier protein (ACP) synthase III] converts malonyl-ACP to acetoacyl-ACP in bacterial fatty acid biosynthesis and multiple studies have reported the essentiality of FabH.^20,21^ Several FabH inhibitors have been reported with potent activity against MRSA.^4,22–24^ Among those, Oxa2 was found to bind with FabH Cys112 which is the same cysteine identified in our proteomic result. Oxa2 showed a MIC range from 0.25∼0.5 µg/mL (1∼2 µM) against a panel of *S. aureus*. However, Oxa2 is not active against gram-negative strains while 10-F05 is active against gram-negative strains.

We also follow up with other cysteine targets of 10-F05, MiaA Cys273 and PdxY Cys121. These two cysteines have slightly lower ligandability and they were also engaged by 10-L07 (**Figure 4C**). MiaA generates the i^6^A-37 tRNA by catalyzing the addition of an isopentenyl group onto the N6-nitrogen of Ade-37 next to the anticodon. The product is subsequently methylthiolated by the radical-S-adenosylmethionine enzyme MiaB to yield ms^2^i^6^A-37.^25^ This modification is known to enhance the interaction of tRNA with UNN codons (Phe, Leu, Ser, Tyr, Cys, Trp), thus promoting reading frame maintenance and translational fidelity.^26,27^ MiaA is necessary for bacterial fitness and virulence in diverse host niches. Intraperitoneal injection of *miaA* KO extraintestinal pathogenic *E. coli* showed significantly lower survival rates compared to the WT *E. coli* strain in a mouse model.^25^ Given its well-defined role in regulating bacterial virulence, MiaA appears to be a novel antibiotic target. Despite MiaA’s conservation in prokaryotes, the identified Cys273 is only conserved in some bacteria strains and its function remains unclear (**Figure S9**).

The third hit PdxY, is known as a pyridoxal kinase involved in the pyridoxal 5′-phosphate (PLP) salvage pathway. However this activity is significantly lower compared to PdxK, indicating that its physiological function remains unknown.^28,29^ PdxY Cys121 is involved in substrate/pyridoxal binding through covalent interaction.^30^

### Biochemical and genetic validation of FabH, MiaA, and PdxY as relevant targets for 10-F05

To confirm the interaction between 10-F05 and the three potential target proteins identified by proteomics, we expressed and purified recombinant Sf_FabH, Sf_MiaA and Sf_PdxY proteins from *E. coli* BL21 strains and visualized the cysteine labeling using tetramethylrhodamine-5-iodoacetamide (5-TMRIA), with competition from 10-F05. As expected, preincubation of 10-F05 decreased the 5-TMRIA labeling in a dose-dependent manner (**Figure 5A**). Time-dependent labeling revealed rapid covalent bond formation between 10-F05 and Sf_FabH (**Figure 5B**). Furthermore, 10-F05 was able to bind with *S. aureus* FabH (Sa_FabH, **Figure 5C**), which shared 39.9% sequence similarity with Sf_FabH. FabH is highly conserved among (**Figure 5D**) while MiaA Cys273 and PdxY are not highly conserved in different bacteria (**Figure S9**), suggesting that FabH is an important target for 10-F05. To test this, we obtained a reported FabH inhibitor Platencin (**Figure 5E, left**) and measured the MIC in the 10-F05 resistant *S. aureus* strain compared to WT strain.^31^ As expected, the 10-F05 resistant *S. aureus* also exhibited resistance to Platencin (**Figure 5E, right**).

**Figure 5.**
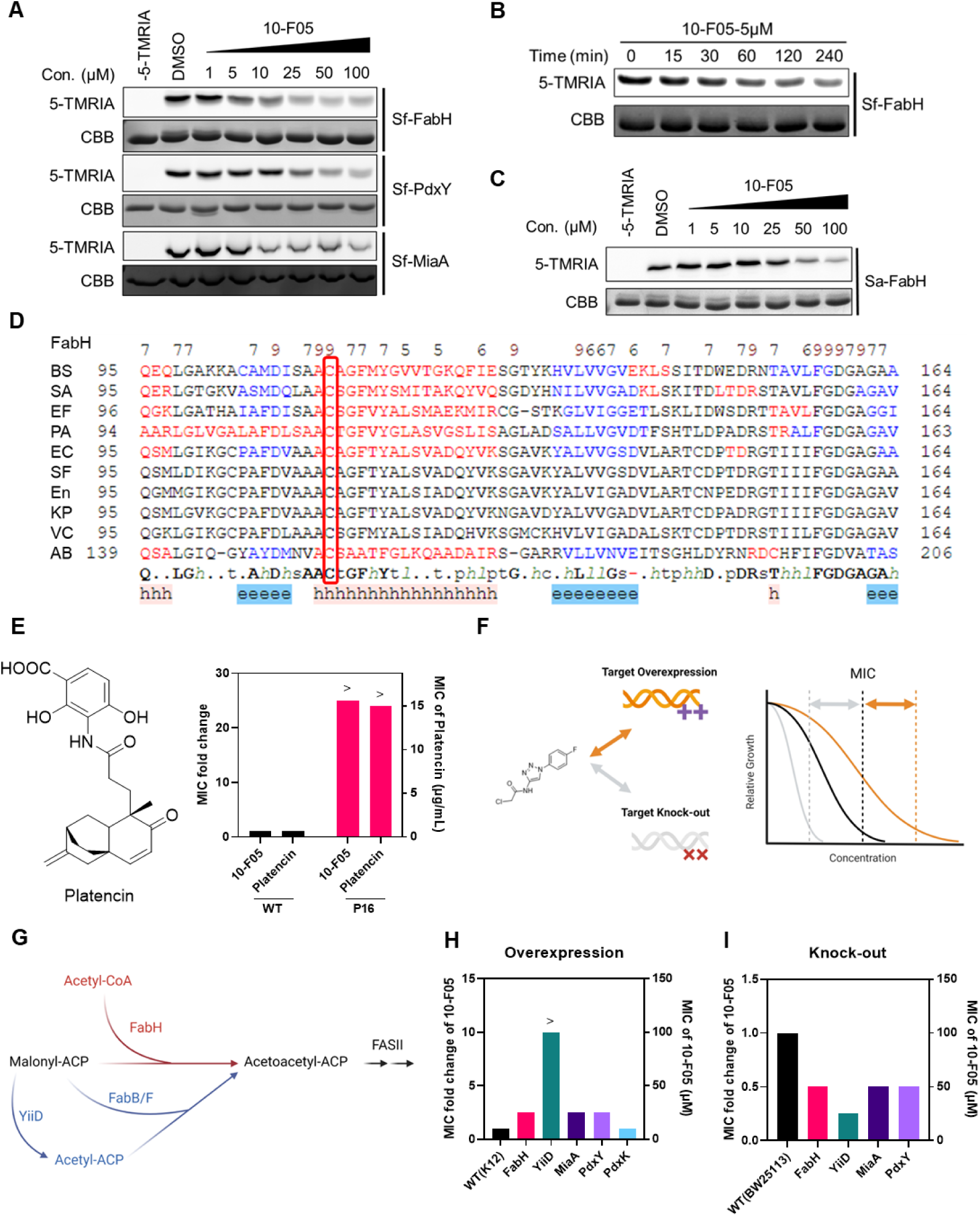
Biochemical and chemogenetic validations suggested that FabH is the major target of 10-F05 mediated growth inhibition. (**A**) In-gel fluorescence labeling of three purified *S. flexneri* target proteins. Purified proteins were incubated with 10-F05 at indicated concentrations for 1 hour and labeled by 5-TMRIA to visualize the unbound cysteines. Uncropped gel images are shown in Supporting Information. (**B**) Time dependent in-gel fluorescence labeling of *S. flexneri* FabH protein. Purified proteins were incubated with 5 µM of 10-F05 for indicated time and labeled by 5-TMRIA to visualize the unbound cysteines. (**C**) In-gel fluorescence labeling of *S. aureus* FabH protein. (**D**) Structural-based sequence alignment of FabH proteins from different bacteria species. BS: *B. subtilis* strain 168; SA: *S. aureus* MSSA476; EF: *E. faecium*; PA: *P. aeruginosa* ATCC 15692; EC: *E. coli* K12; SF: *S. flexneri* M90T; En: Enterobacter sp. (strain 638); KP: *K. pneumoniae* subsp. pneumoniae ATCC 700721; VC: *V. cholerae* serotype O1; AB: *A. baumannii* NCGM 237. (**E**) Left, structure of a reported FabH inhibitor, Platencin. Right, MIC fold change of 10-F05 and Platencin tested in WT and 10-F05 resistant *S. aureus* (Passage 16 from Figure 3B). >: No full growth inhibition was observed at the highest concentration tested, indicating that the real MIC was higher than the number reported here. (**F**) Scheme for chemical genetic interaction. MIC shifts correspond to the target overexpression and knock-out. (**G**) YiiD bypasses FabH to generate acetoacetyl-ACP in *E. coli*. (**H**) MIC fold change of 10-F05 tested in target gene overexpression *E. coli* strains (ASKA collection). (**I**) MIC fold change of 10-F05 tested in target gene knock out *E. coli* strains (Keio collection).

To further confirm the identified protein targets of 10-F05 in bacteria, we used over-expression and knock-out (KO) *E. coli* strains (**Figure 5F**). We validated the ASKA strains over-expressing the selected proteins using western blot analysis (**Figure S10**).^32^ We then measured the MIC of 10-F05 in these strains compared to the parental WT strain (**Figure 5H**). Over-expressing FabH resulted in 2.5-fold increase the MIC. Overexpression of YiiD (also known as FabY), which compensates for the loss of FabH (**Figure 5G**),^33,34^ fully rescued the growth inhibition by 10-F05. This difference in tolerance between FabH and FabY over-expression could be attributed to the fact that the over-expressed FabH is still inhibited by excess 10-F05, but the over-expressed YiiD is not inhibited by 10-F05. Similarly, the MIC of 10-F05 increased 2.5-fold in MiaA and PdxY over-expressing *E. coli* strains. Interestingly, over-expressing PdxK had no effect on 10-F05 treatment, suggesting that PdxY likely does not function in pyridoxal salvage. The MIC increase caused by FabH/MiaA/PdxY gene over-expression further supports that they are relevant target proteins of 10-F05.

We obtained the knock-out strains from Keio collection^35^ and compared the MIC difference of 10-F05 in WT and *fabH/miaA/pdxY* KO strains. Knocking out any of the three targeted genes sensitized *E. coli* to 10-F05 treatment, resulting in a 2-fold MIC change (**Figure 5I**). Notably, the *yiiD* KO strain showed the largest MIC change consistent with the overexpression data. Thus, our chemical proteomics combined with the biochemical and genetic data revealed that multiple targets contribute to the observed antimicrobial activity of 10-F05, but given the strongest effect of yiiD (which bypasses FabH) overexpression or deletion on the sensitivity to 10-F05, FabH seems to be the major target underlying the antibacterial activity of 10-F05.

### 10-F05 disrupts MiaA and tRNA substrate binding *in vitro*

Among the top three cysteine targets we identified, the function of MiaA Cys273 remains unclear. Thus, we aimed to investigate the impact of covalent modification of MiaA Cys273 by 10-F05 on MiaA’s function. Sf_MiaA contains a total of two cysteines. To confirm that 10-F05 reacts with MiaA Cys273, we purified the MiaA C273A mutant and performed 5-TMRIA labeling. The C273A mutant had decreased 5-TMRIA labeling and the signal was not decreased by 10-F05 competition, indicating 10-F05 specifically reacts with MiaA Cys273, consistent with our proteomic data (**Figure 6A**).

**Figure 6.**
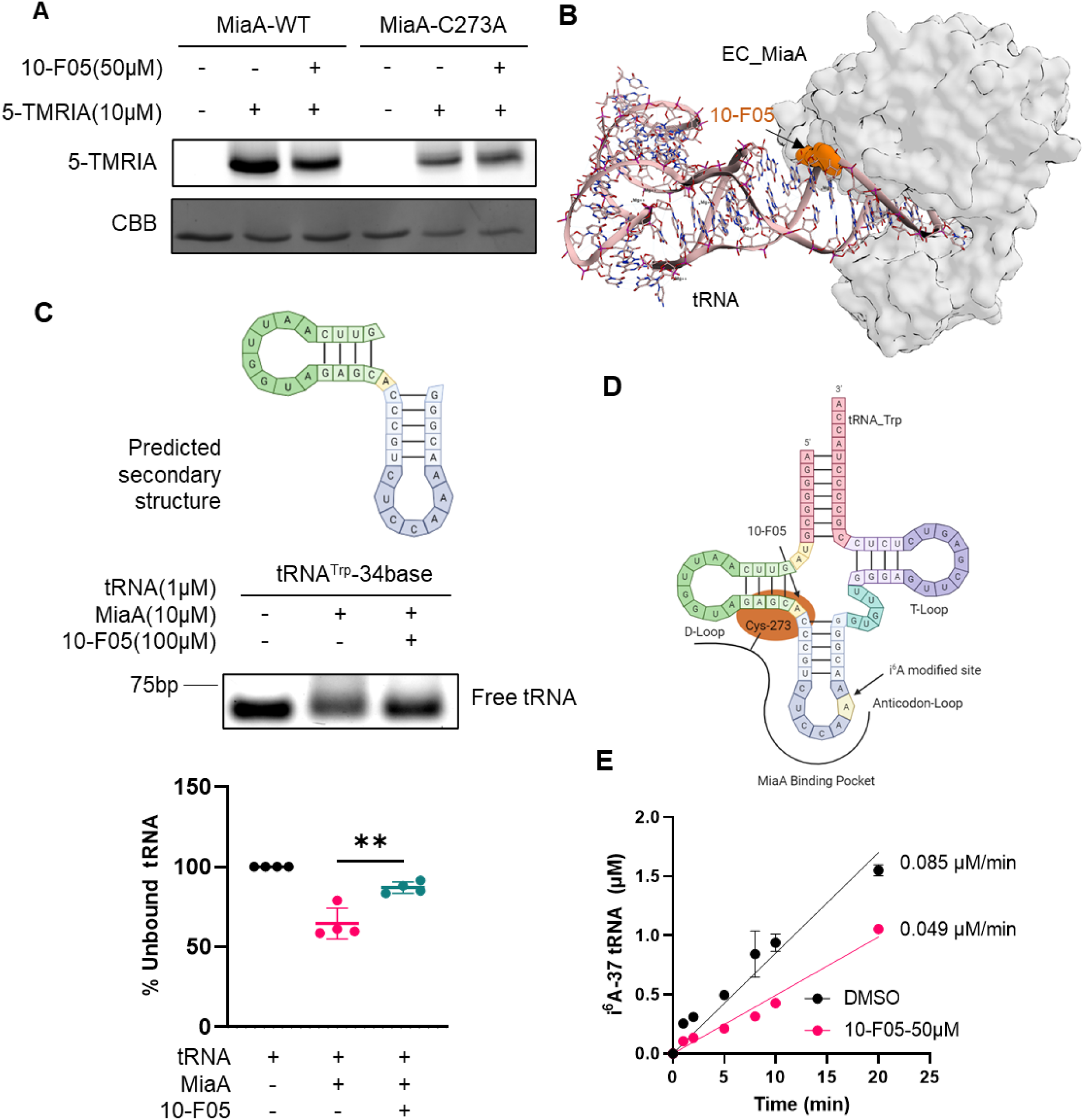
10-F05 disrupts MiaA and tRNA substrate binding. (**A**) In-gel fluorescence of purified WT and C273A MiaA with and without 10-F05 as the competitor. Uncropped gel images are shown in Supporting Information. (**B**) Computational model of 10-F05 bound to *E. coli* MiaA (PDB: 2ZM5) by MOE (2020). 10-F05 is shown as orange surface. The 10-F05 binding site overlaps with the tRNA substrate binding site. (**C**) Top: predicted secondary structure of designed tRNA sequence; Middle: gel-based tRNA mobility shift assay; Bottom: quantification of the ratio of unbound (free) tRNA (N=4). (**D**) Simplified model showing that the linker region between anticodon loop and D loop of tRNA is blocked by 10-F05 addition. (**E**) i^6^A-37 tRNA formation rate monitored by LC-MS in an *in vitro* enzymatic assay (N=2). The p values of unpaired two-tailed t-tests were calculated in GraphPad Prism (version 9.3.1). **p < 0.01.

MiaA Cys273 is located near the tRNA binding site (**Figure S11A**). Covalent docking was performed to visualize the interaction of 10-F05 and MiaA (**Figure S11B**). The predicted covalent docking structure revealed a spatial overlap between 10-F05 and the tRNA substrate, suggesting that the covalent modification by 10-F05 could interfere with tRNA binding (**Figure 6B**, 10-F05 is shown in orange surface). To test this, we designed a 34-base tryptophan tRNA (tRNA^Trp^_34) sequence predicted to mimic the tRNA D-loop and anticodon loop,^36^ and conducted a gel mobility shift assay (**Figure 6C**). We first incubated the purified MiaA protein with 10-F05 or DMSO, followed by the addition of the tRNA substrate. Bound and unbound tRNA substrate were separated using electrophoresis and visualized using Gel-Red staining, which were further quantified to calculate the unbound tRNA ratio. The unbound tRNA^Trp^_34 ratio increased from 64% to 87% after preincubation with 10-F05, in agreement with our hypothesis (**Figure 6C** and **Figure 6D**). As expected, the MiaA C273A mutant, which does not react with 10-F05, was not affected much by 10-F05 treatment in tRNA binding (**Figure S12**).

To examine whether the substrate blocking effect of 10-F05 could influence MiaA’s activity, we performed an *in vitro* enzymatic activity assay.^37^ The reaction kinetics were measured by monitoring the i^6^A-37 tRNA product using LC-MS. As expected, preincubation of 10-F05 with MiaA decreased the initial velocity of modified tRNA forming rate by 1.7-fold using tRNA^Trp^_34 as the substrate (0.085 *vs* 0.049 µM/min, **Figure 6E**).

### 10-F05 treatment increases translation error and decreases stress resistance through MiaA inhibition

We next investigated whether 10-F05 inhibits MiaA function in bacteria (**Figure 7A**). We utilized a dual-luciferase reporter plasmid to quantify translation fidelity change in *miaA* KO *E. coli* strains^25^. Our results revealed an elevated translation error in the *miaA* KO strain compared to the WT strain, which aligns with previously reported results.^25^ Subsequently, we treated both the WT and *miaA* KO strains with 5 µM of 10-F05. The results indicated that 10-F05 increased the translation error in the WT strain but not in the *miaA* KO strain. This observation strongly suggests that 10-F05 reduces translation fidelity by inhibiting the activity of MiaA (**Figure 7B)**. Interestingly, the measured translation error in 10-F05-treated WT *E. coli* cells was even higher than that in the *miaA* KO *E. coli* cells, likely because the *miaA* KO cells were somehow adapted to the increased translation error in the absence of i^6^A modified tRNA.

**Figure 7.**
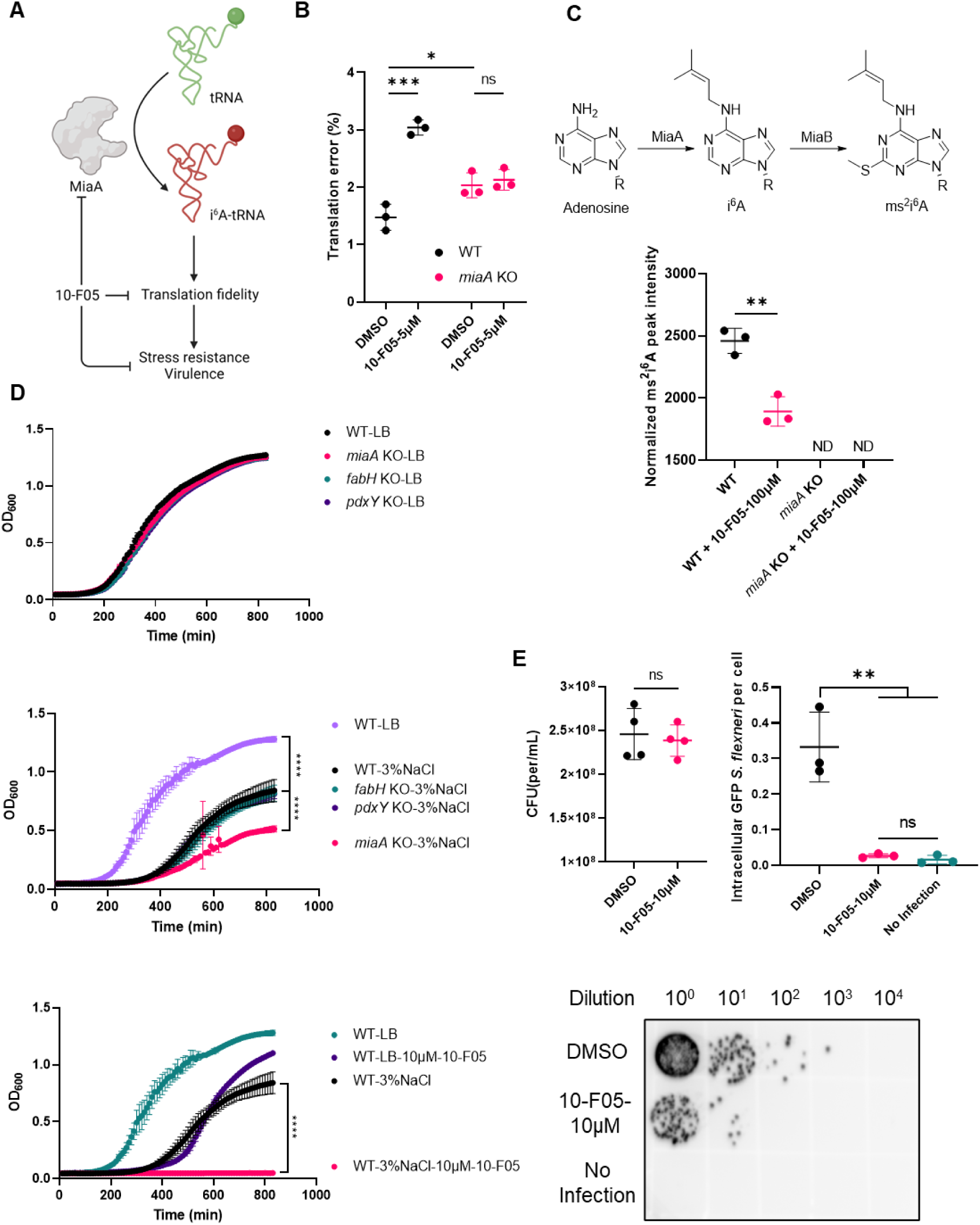
10-F05 increases translation error through MiaA inhibition *in vivo* to decrease bacterial stress resistance and virulence. (**A**) Scheme showing MiaA’s role in regulating bacteria stress resistance and virulence. (**B**) Translation error measured using -1 frame shift reporter plasmids transformed in WT (BW25113) and *miaA* KO *E. coli* strains. (**C**) Top, biosynthesis pathway of i^6^A and ms^2^i^6^A tRNA. Bottom, normalized ms^2^i^6^A peak quantified by LC-MS. Corresponding tRNA samples were extracted from WT and miaA KO *E. coli* strains with or without 100 µM of 10-F05 treatment. ND: not detected. (N=3). (**D**) Top, growth curves of *E. coli* cultured in LB media. Middle, growth curves of *E. coli* cultured in 3% NaCl-LB media. Bottom, growth curve of WT *E. coli* cultured with or without 10 µM of 10-F05 in LB and 3% NaCl-LB media. (**E**) Top left, comparison of colony forming units (CFU) difference after 1-hour treatment with 10 µM of 10-F05 (N=4). Top right, comparison of intracellular GFP-*S. flexneri* level using Cytation 5 imaging. Intracellular GFP foci was quantified using ImageJ (1.53t). Three images of each group were acquired and analyzed. Bottom, intracellular *S. flexneri* level quantified using growth on agar plate. The p values of unpaired two-tailed t-tests were performed in GraphPad Prism (version 9.3.1). *p < 0.05, **p < 0.01, ***p < 0.001, ****p < 0.0001, and ns, not significant.

To further validate that 10-F05 inhibits MiaA in cells, we proceeded to purify the modified tRNA from *E. coli* and enzymatically digested it into individual ribonucleosides using established methods.^38,39^ Subsequently, we quantified the presence of i^6^A and ms^2^i^6^A (from i^6^A by the action of MiaB) using LC-MS analysis (**Figure 7C, top**). The i^6^A modification level was extremely low and below the detectable limit of our LC-MS instrument. Nonetheless, we were able to detect a distinct peak corresponding to ms^2^i^6^A (same retention time as the synthetic standard of ms^2^i^6^A, **Figure S13**). Importantly, this peak was only observed in the WT strain but not in the *miaA* KO strain, confirming the absence of i^6^A modification in the latter. As expected, when we treated the *E. coli* strain (starting from ∼10^8^ inoculation) with 100 µM of 10-F05 for 3 hours, we observed a ∼24% decrease in the level of ms^2^i^6^A compared to the DMSO control group (**Figure 7C, bottom** and **Figure S13**). This reduction in ms^2^i^6^A level further supports that 10-F05 inhibits the activity of MiaA in the bacteria.

Previous studies conducted in *S. flexneri* and *E. coli* demonstrated that the deletion of *miaA* led to a reduction in the expression of virulence factors and altered stress resistance (**Figure 7A)**.^25,40,41^ Thus, we tested whether the pharmacological inhibition of MiaA by 10-F05 causes similar effects. The fabH, miaA, or pdxY KO strains exhibited no significant difference in growth compared to the WT strain when cultured in LB media (**Figure 7D, top**). However, the deletion of MiaA resulted in decreased *E. coli* growth rate under osmotic stress (**Figure 7D, middle**). The addition of sub-MIC concentration of 10-F05 completely shut down WT *E. coli* growth in 3% NaCl LB media but not in normal LB media, indicating that pharmacological inhibition of MiaA sensitized bacteria to osmotic stress (**Figure 7D, bottom**).

To evaluate the impact of MiaA inhibition on bacterial virulence, we pre-treated GFP-tagged *S. flexneri* M90T (starting from ∼ 3.2 × 10^8^ inoculation) with 10-F05 for 1 h and then infected Hela cells. *S. flexneri* is an intracellular pathogen and its pathogenesis requires entering the host cells. We assessed changes in virulence by quantifying the number of intracellular bacteria. No growth inhibition, as indicated by the colony-forming ability, was observed after 1-hour treatment with 10 µM of 10-F05 (**Figure 7E, top left**). However, the average number of intracellular *S. flexneri* was significantly reduced following 10-F05 treatment (**Figure 7E, top right** and **Figure S14**). Similarly, an agar-based quantification demonstrated a significant decrease in the intracellular level of *S. flexneri*, indicating decreased virulence after 10-F05 treatment (> 100-fold change, **Figure 7E, bottom**). Overall, our findings revealed that the pharmacological inhibition of MiaA leads to a similar phenotype as *miaA* KO in bacterial pathogens, suggesting MiaA as a promising target that controls bacterial virulence and stress resistance.^25,40^

## Discussion

By screening a library of ∼3,200 cysteine-targeting compounds, we discovered a series of aromatic ring-conjugated chloromethyl ketone scaffolds exhibiting potent antibacterial activity. The leading compound, 10-F05, showed broad-spectrum activity and slow resistance development rate. The competitive ABPP workflow enabled us to quickly identify the molecular targets (FabH, MiaA, and PdxY) of 10-F05. We further validated their physiological relevance in mediating the growth inhibitory activity of 10-F05 through chemical genetic interactions.

FabH has previously been indicated as an antibiotics target. Various chemical scaffolds, including 1,3,5-oxadiazin-2-one,^22^ oxadiazolones,^4^ platencin and its derivatives,^23,31^ and benzoylaminobenzoic acid,^42^ have been reported as inhibitors of FabH, through covalent or non-covalent interactions. However, despite their distinct structures, these FabH inhibitors have shown a loss of antibacterial activity in gram-negative bacterial strains due to effective drug efflux.^24,42^ The compound discovered here, 10-F05, is able to inhibit FabH in gram-negative bacteria. It is possible that its smaller size leads to better cell permeability, or its rapid target engagement may lead to a reduction in excretion levels through TolC, thereby retaining its potency against gram-negative bacteria.

Notably, MiaA and PdxY have not been recognized as antibiotics targets and no inhibitors targeting them have been reported to date. Therefore, 10-F05 represents a promising starting scaffold for the development of selective inhibitors for these new targets. Moreover, we validated that MiaA is relevant antibacterial target for 10-F05. Through examining published MiaA structures and experimental validation, we discovered that the covalent addition of 10-F05 to MiaA Cys273 disrupts its interactions with the tRNA substrates, inhibiting MiaA function and leading to reduced stress resistance and virulence. The 10-F05 MIC changes observed in strains overexpressing PdxY and PdxK further suggests an unidentified role for PdxY that is unrelated to PLP salvage.

Reactivity is a crucial factor to consider in covalent drug design as it influences off-target effects. Traditional covalent drugs often employ weak electrophilic warheads to facilitate optimized covalent bond formation primarily within the desired binding complex. This minimizes off-target protein labeling. Our study here showed that many commonly used warhead scaffolds lack antibacterial activity against gram-negative bacterial strains. This could be attributed to their low affinity, as most compounds in the library are fragment-like, resulting in a low secondary second-order rate constant of inactivation (*kinact*/*KI*). In contrast, the highly reactive aromatic ring-conjugated chloromethyl ketones likely increases the chances of engagement of these fragment-like moieties.

We initially had concerns about the promiscuity of 10-F05 due to its high reactivity. However, our proteomic results demonstrated reasonable selectivity of 10-F05 in live bacterial cells. Furthermore, a previous study reported that promiscuity does not necessarily correlate with reactivity.^16^ Future experiments are necessary to explore the mechanism underlying the target selectivity of 10-F05. We also note that the ability of 10-F05 to target several bacterial proteins has a potential advantage - the bacteria are less likely to develop resistance to 10-F05 (Figure 3) compared to other common antibiotics tested (kanamycin and methicillin). This is because it is difficult for bacteria to accumulate mutations in all of the target proteins.

It is also important to note that the high reactivity of cysteine covalent drugs can result in a decreased stability for *in vivo* use. This is due to their rapid consumption by glutathione, which may explain the lower cytotoxicity of chloromethyl ketone scaffolds compared to chloroacetamide in mammalian cells. These findings highlight the complex role of reactivity that needs to be considered in covalent antibiotic design. Further optimization of 10-F05 is required for developing clinical antibiotic candidates. Additionally, it is crucial to develop a more diversified covalent library with a broad range of reactivity to effectively target bacterial cysteines with various properties.

## Conclusion

Our study demonstrates that combining covalent compound library phenotypic screening (bacterial growth) with ABPP is an efficient way to identify new protein targets (such as MiaA) as well as lead compounds. Traditional non-covalent compound library phenotypic screening is limited by the difficulty in identifying the protein targets of the hit compounds. The use of covalent compound library and ABBP chemical proteomics effectively overcomes this limitation. This approach therefore has the potential to open new research directions in drug discovery and chemical biology.

## Acknowledgement

We thank Dr. Matthew A. Mulvey and Alexis Rousek from The University of Utah (pCWR44, pCWR45) for their gift of plasmids. We thank the National BioResource Project (NRBP) of Japan for providing the E. coli Keio and ASKA strains. We thank Dr. Tobias Dörr from Cornell University for providing *V. cholerae* SAD30. We thank Dr. Neal M. Alto from UT Southwestern for providing *S. flexneri* 5a M90T. We thank Drs. Sheng Zhang, Qin Fu, and the Cornell Proteomics Facility for helping with the proteomics studies. We thank HHMI Transformative Technology 2019 Program for the purchase of the Orbitrap Eclipse mass spectrometer.

