## Supporting Information for "Phenotypic screening of covalent compound libraries identifies chloromethyl ketone antibiotics and MiaA as a new target"

### Supplementary Tables

**Table S1.** Hit count with different warhead scaffolds.

| Warhead | Total amount in Cys-library | Hit amount (SA) | Hit amount (VC) |
|---|---|---|---|
| 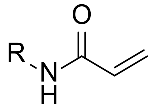 | 960 | 1 | 1 |
| 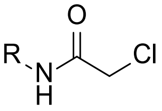 | 751 | 11 | 0 |
| 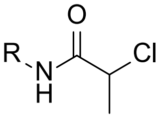 | 529 | 0 | 0 |
| 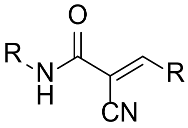 | 640 | 4 | 1 |
| 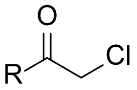 | 210 | 31 | 15 |
| 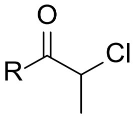 | 26 | 0 | 0 |
| 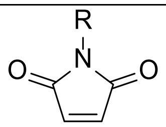 | 56 | 1 | 0 |
| 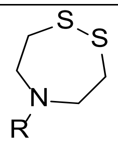 | 25 | 0 | 0 |
| 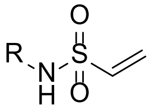 | 1 | 0 | 0 |
| 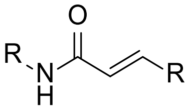 | 2 | 0 | 0 |

SA: *S. aureus*, VC: *V. cholerae*

**Table S2.** MIC table of negative control compounds.

| Structure | Compound Name | MIC (μM) | | | |
|---|---|---|---|---|---|
| | | ***S. flexneri*** | ***V. cholerae*** | ***E. coli*** | ***S. aureus*** |
| 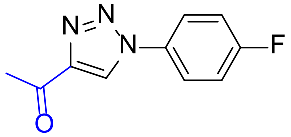 | 10-F05-N | >100 | >100 | >100 | >100 |
| 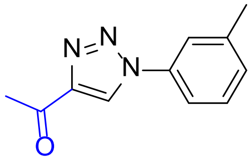 | 10-L07-N | >100 | >100 | >100 | >100 |
| 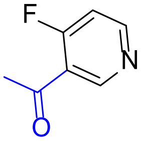 | 10-I09-N | >100 | >100 | >100 | >100 |
| 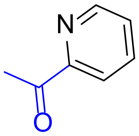 | 10-J03-N | >100 | >100 | >100 | >100 |

### Supplementary Figures

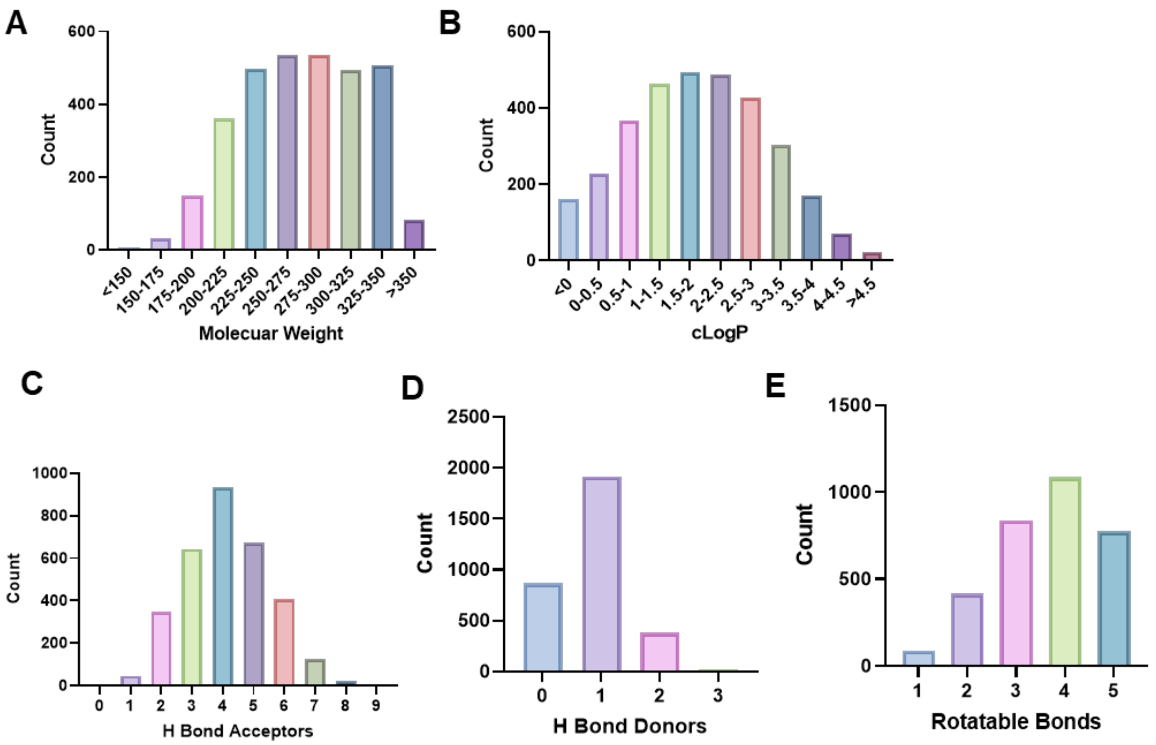

**Figure S1.** Molecular properties of the Cys-library used. (**A**) Molecular weight. (**B**) cLogP. (**C**) H bond acceptors. (**D**) H bond donors. (**E**) Rotatable bonds. All the properties were calculated using Data Warrior.

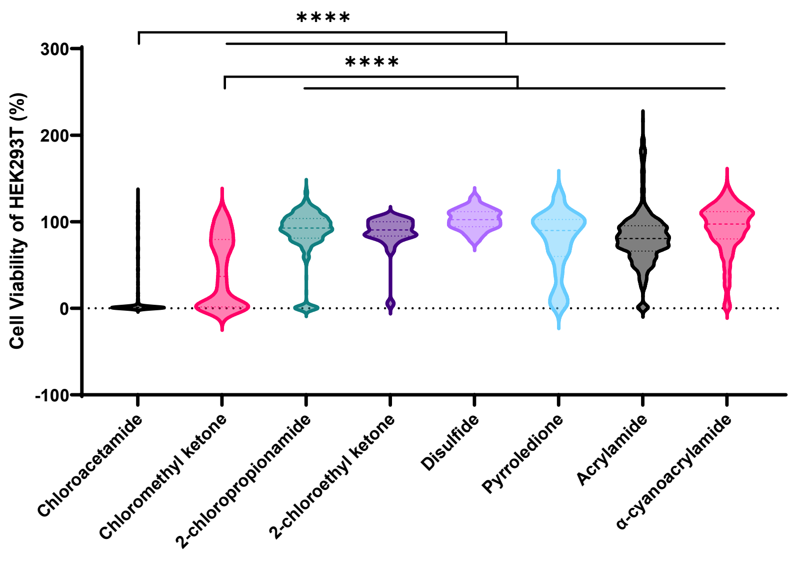

**Figure S2.** Cytotoxicity profiling of the Cys-library at 25 µM. Cell viability of HEK293T cells was measured using the CellTiter-Glo 2.0 cell viability assay kit after a two-day incubation period with the Cys-library. Each group was compared to chloroacetamide and chloromethyl ketone scaffolds using unpaired two-tailed t-tests. **** P <0.0001.

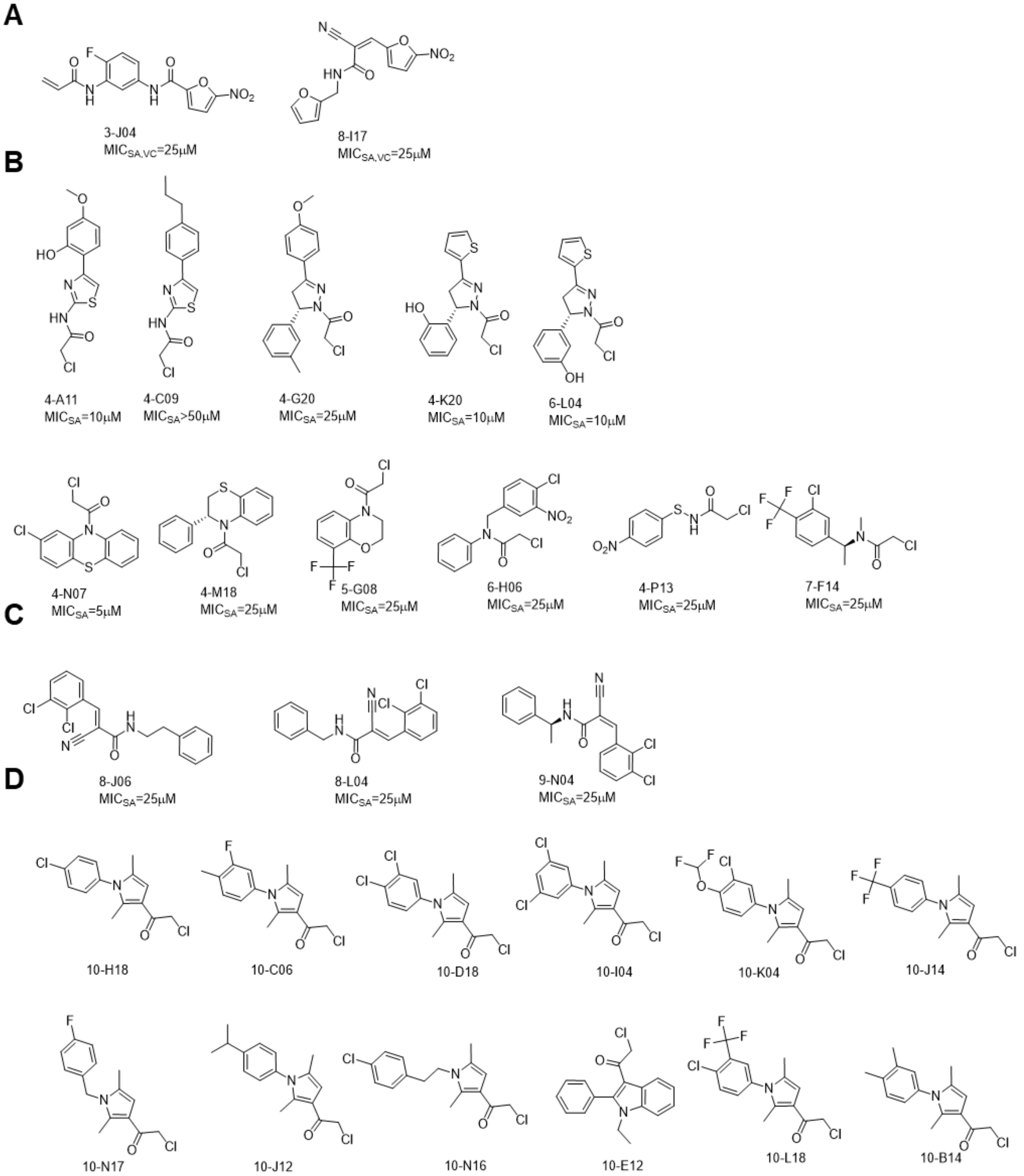

**Figure S3.** Compounds that were filtered out due to high cytotoxicity in HEK293T. (**A**) Nitrofuran containing compounds. (**B**) Chloroacetamides. (**C**) α-Cyanoacrylamides. (**D**) Chloromethyl ketones.

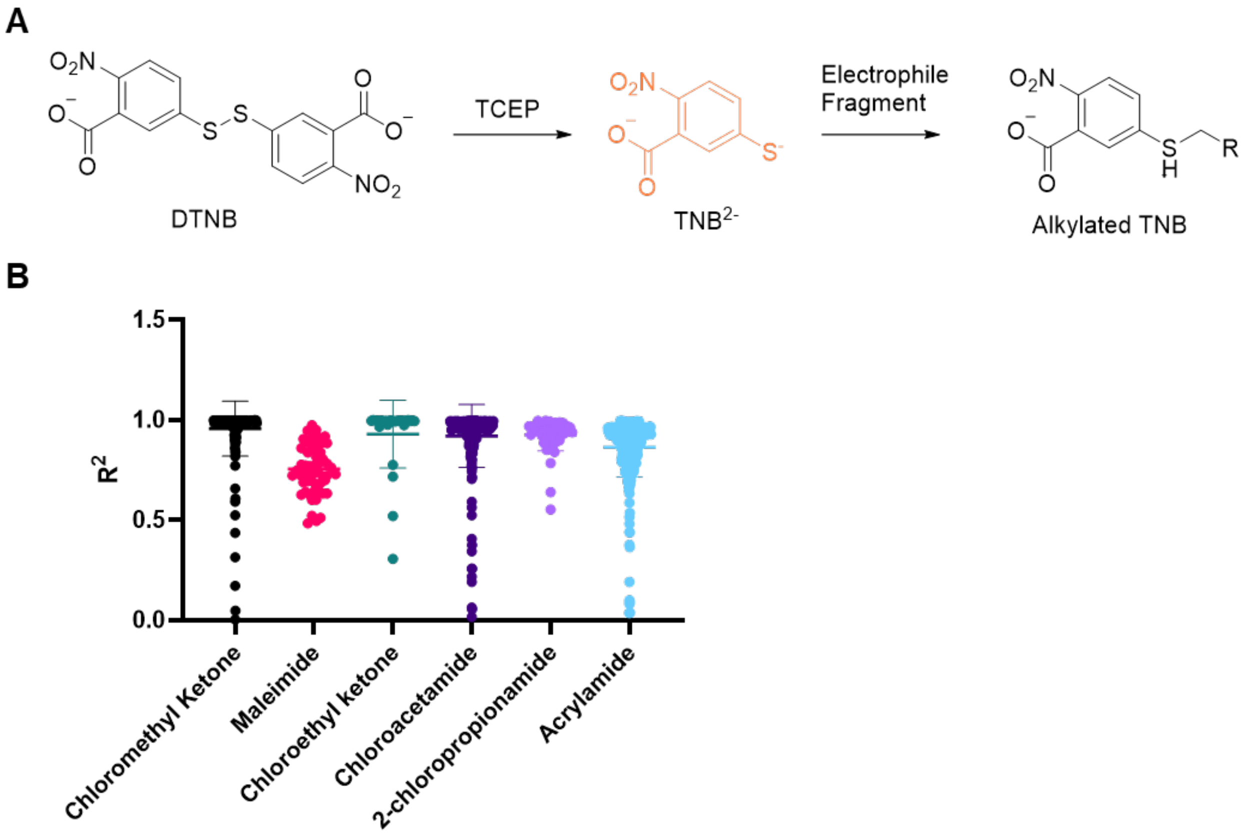

**Figure S4.** DTNB reactivity assay. (**A**) Mechanism of reduced DTNB assay. (**B**) Comparison of R square from linear regression clustered by different warhead scaffolds.

A

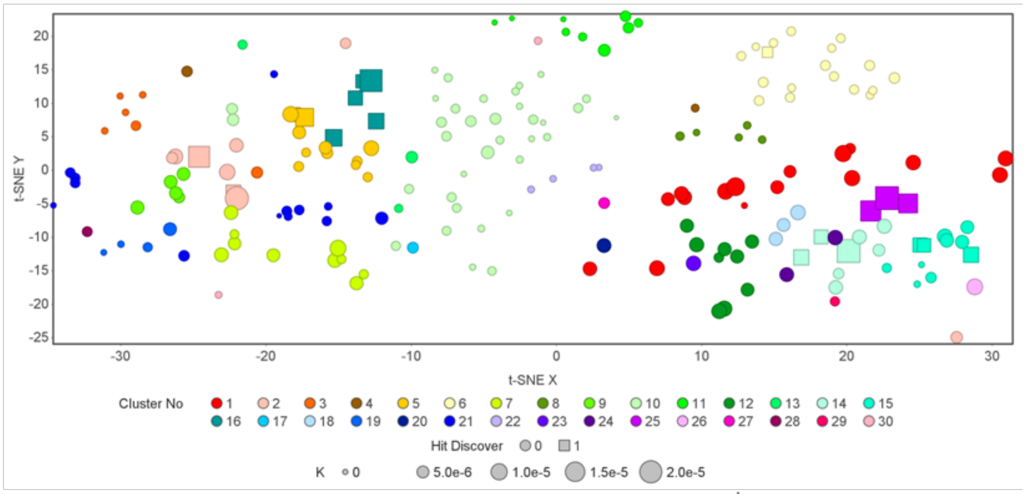
t-SNE Y
t-SNE X
Cluster No 1 2 3 4 5 6 7 8 9 10 11 12 13 14 15 16 17 18 19 20 21 22 23 24 25 26 27 28 29 30
Hit Discover 0 1
K 0 5.0e-6 1.0e-5 1.5e-5 2.0e-5

B

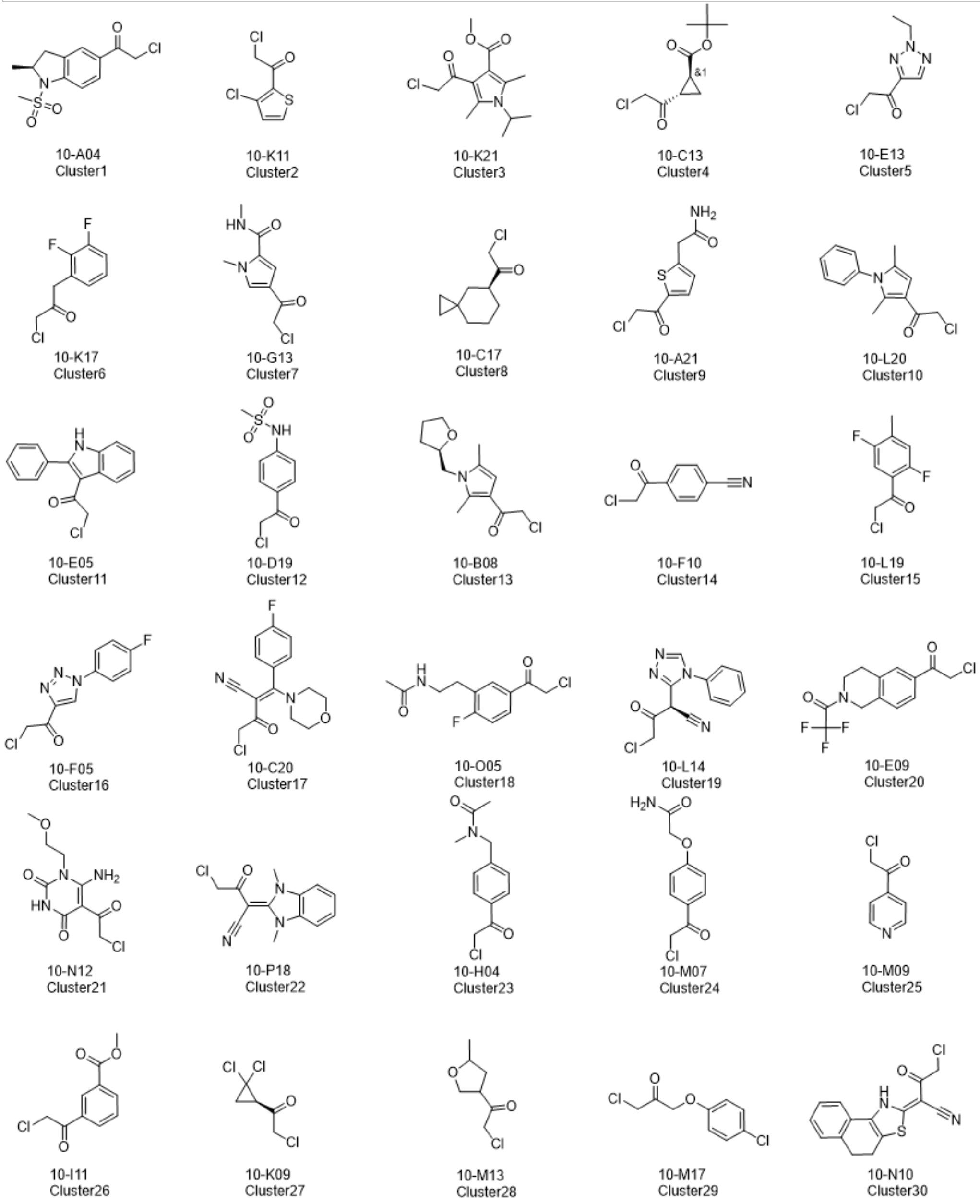
10-A04 Cluster1
10-K11 Cluster2
10-K21 Cluster3
10-C13 Cluster4
10-E13 Cluster5
10-K17 Cluster6
10-G13 Cluster7
10-C17 Cluster8
10-A21 Cluster9
10-L20 Cluster10
10-E05 Cluster11
10-D19 Cluster12
10-B08 Cluster13
10-F10 Cluster14
10-L19 Cluster15
10-F05 Cluster16
10-C20 Cluster17
10-O05 Cluster18
10-L14 Cluster19
10-E09 Cluster20
10-N12 Cluster21
10-P18 Cluster22
10-H04 Cluster23
10-M07 Cluster24
10-M09 Cluster25
10-I11 Cluster26
10-K09 Cluster27
10-M13 Cluster28
10-M17 Cluster29
10-N10 Cluster30

**Figure S5.** SAR analysis of chloromethyl ketone scaffolds. (**A**) Compounds were clustered into 30 clusters labeled with different colors. Representative structures are displayed near each cluster, and the active hits against both *S. aureus* and *V. cholerae* are labeled by squares while others are labeled by circles. The size of the square/circle reflects their reactivity measured by DTNB assay. (**B**) Representative structure of each cluster.

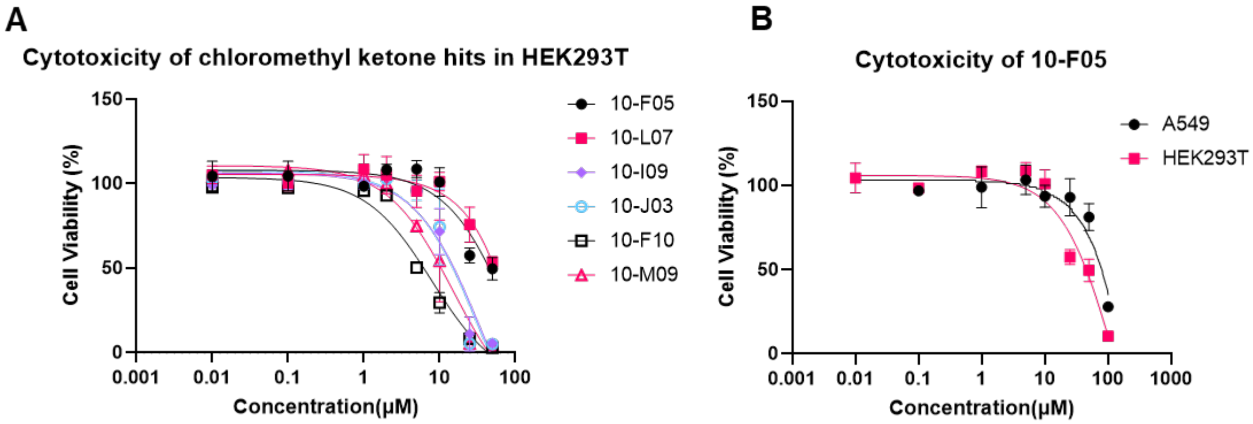

**Figure S6.** (**A**) Cytotoxicity profiling of 10-F05 derivatives in HEK293T cells. (**B**) Cytotoxicity of 10-F05 in A549 and HEK293T cells. Cell viability was assessed using the CellTiter-Glo 2.0 cell viability assay kit after a two-day incubation period.

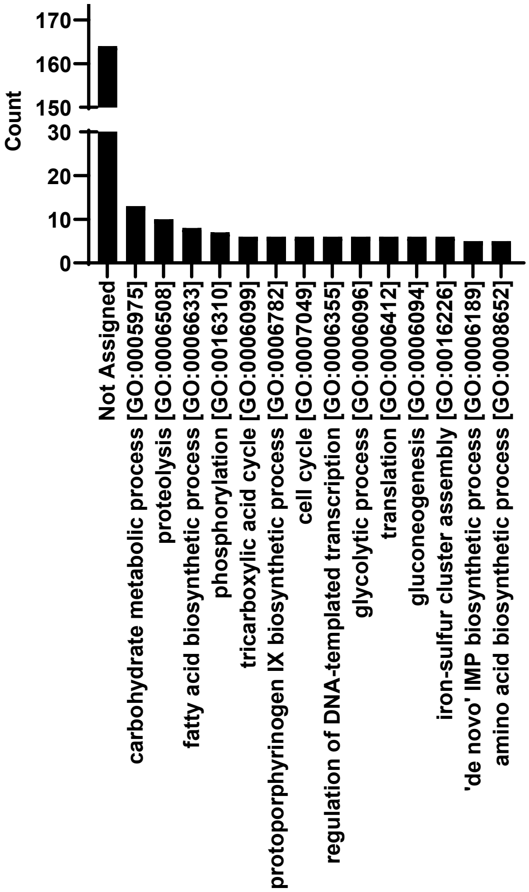

**Figure S7.** Count of quantified protein function for the protein hits identified in the proteomics study. Protein functions were queried from Uniprot Database.

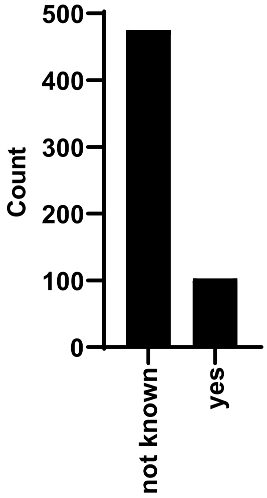

**Figure S8.** Count of quantified essential proteins for the protein hits identified in the proteomics study. Protein essentiality was predicted by NetGene database.

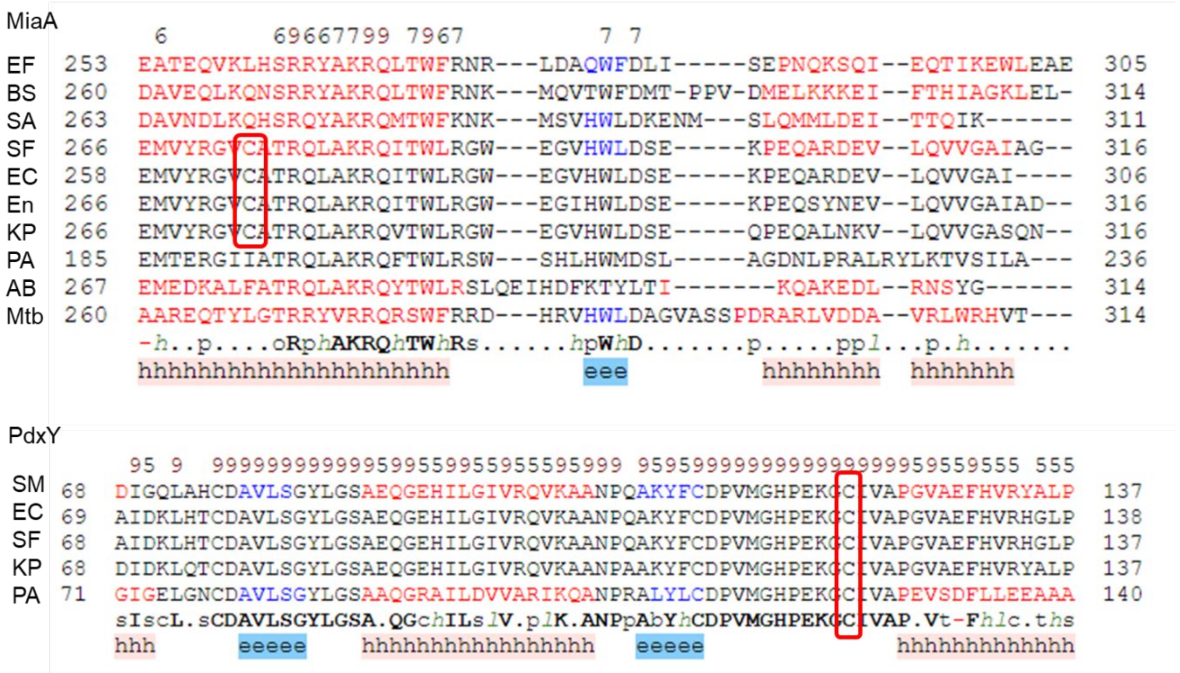

**Figure S9.** Structure-based sequence alignment for MiaA and PdxY. EF: *E. faecium* (AF-A0A7V7GPP3-F1-model_v4); BS: *B. subtilis* strain 168 (AF-O31795-F1-model); SA: *S. aureus* MSSA476 (AF-Q6G9R0-F1-model_v4); SF: *S. flexneri* M90T (AF-A0A4P7TQH3-F1-model_v4); EC: *E. coli* K12 (PDB: 2ZM5); En: *Enterobacter* sp. strain 638 (AF-A4W5R2-F1-model_v4); KP: *K. pneumoniae* strain 342 (AF-B5Y333-F1-model_v4). PA: *P. aeruginosa* ATCC15692 (PDB: 3CRM); AB: *A. baumannii* BAA1790 (AF-A0A059ZUH9-F1-model_v4); Mtb: *M. tuberculosis* ATCC 25177 (AF-A5U679-F1-model_v4). PdxY, SM: *S. typhimurium* strain LT2 (AF-Q8ZPM8-F1-model_v4); EC: *E. coli* K12 (PDB: 1TD2); SF: *S. flexneri* M90T (AF-A0A4P7TT64-F1-model_v4); KP: *K. pneumoniae* IS53 (AF-W1EDE7-F1-model_v4); PA: *P. aeruginosa* PAO1 (PDB: 5B6A). The covalently targeted cysteines are highlighted in red boxes.

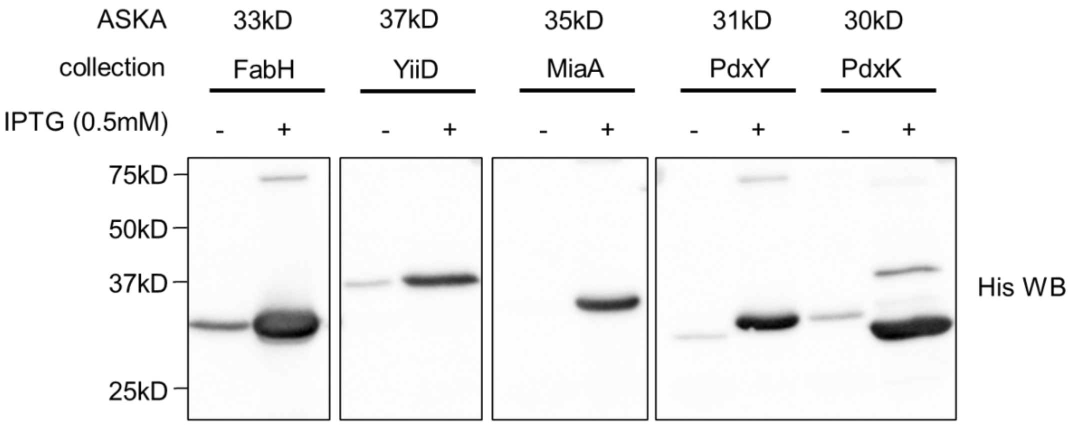

**Figure S10.** Verification of ASKA overexpression strains. IPTG (0.5mM) was added into the cell culture to induce protein overexpression. Protein molecular weight, FabH: 33kD; YiiD: 37kD; MiaA: 35kD; PdxY: 31kD: PdxK: 30kD.

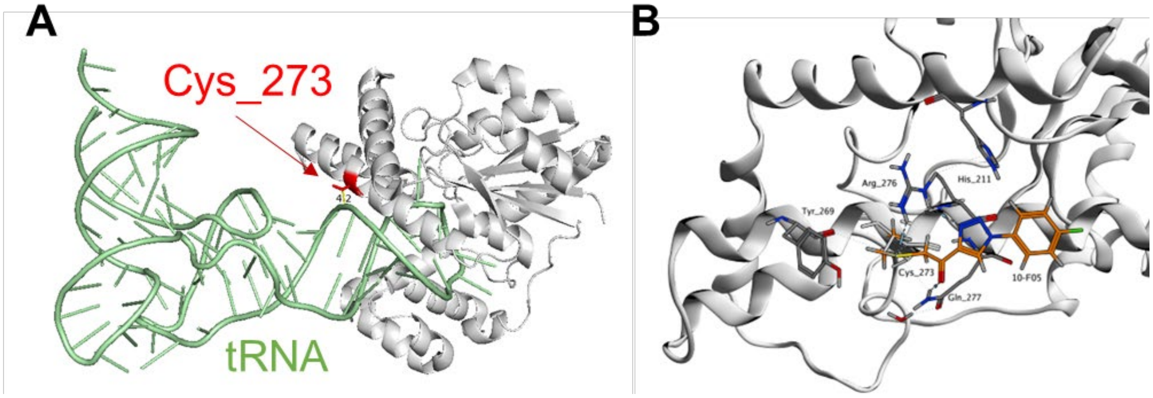

**Figure S11.** (A) MiaA Cys273 is very close to the bound tRNA substrate. *E. coli* MiaA (PDB: 2ZM5). (B) Predicted covalent interaction of 10-F05 with nearby amino acids residues. Hydrogen bonding interactions are indicated by short blue sticks.

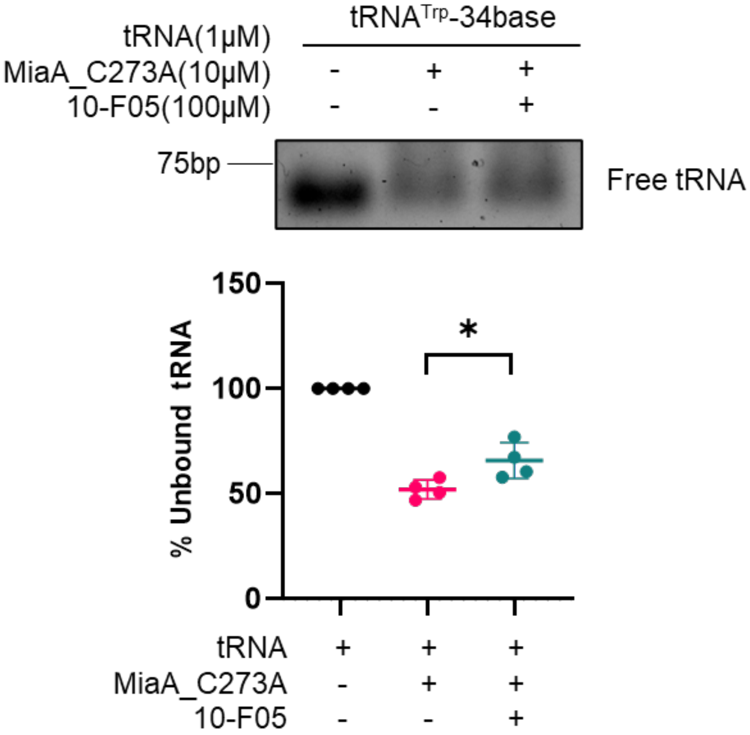

**Figure S12.** 10-F05 only slightly decreases MiaA_C273A binding with tRNA$^{Trp}$-34.

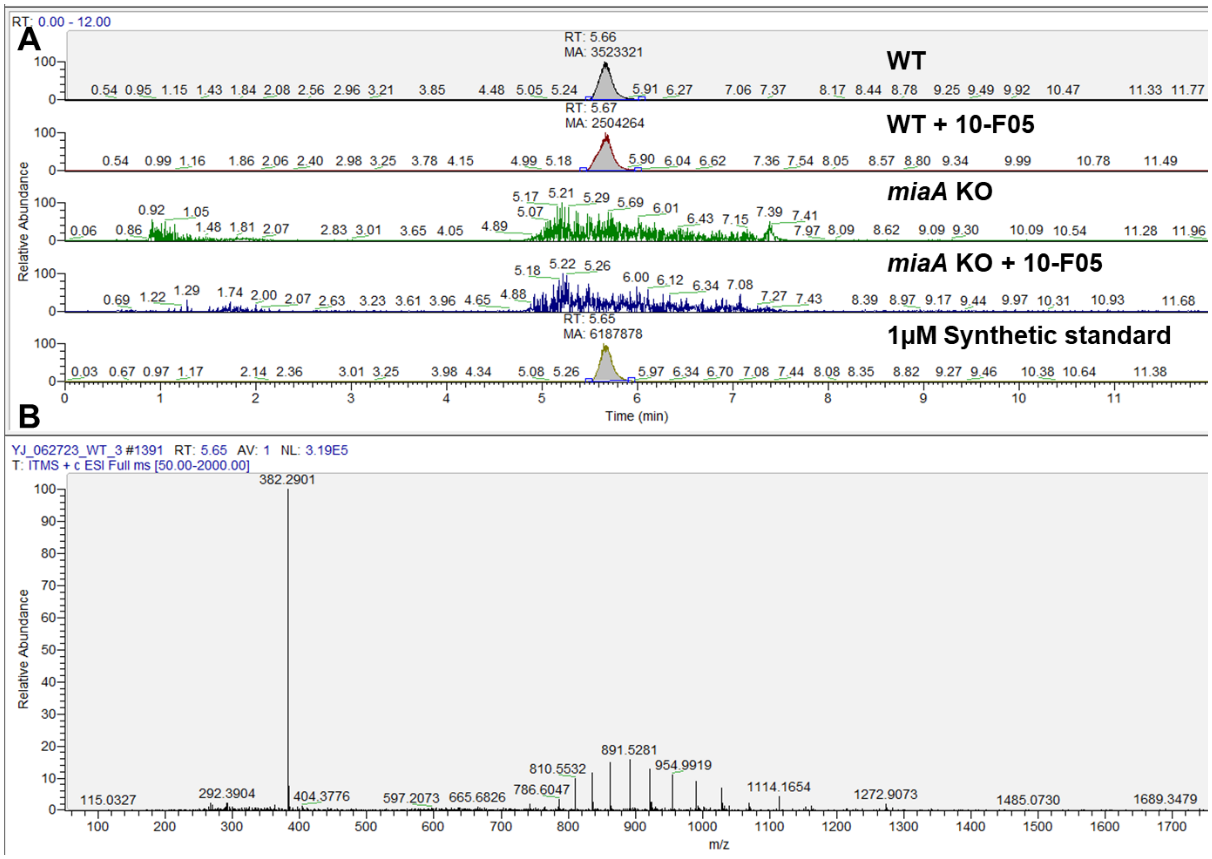

**Figure S13.** (**A**) Representative images of $ms^2i^6A$ peak in LC-MS analysis. Peak corresponding to the $ms^2i^6A$ was searched using ms/z range from 381.5000~382.5000. (**B**) MS spectrum of the $ms^2i^6A$ peak.

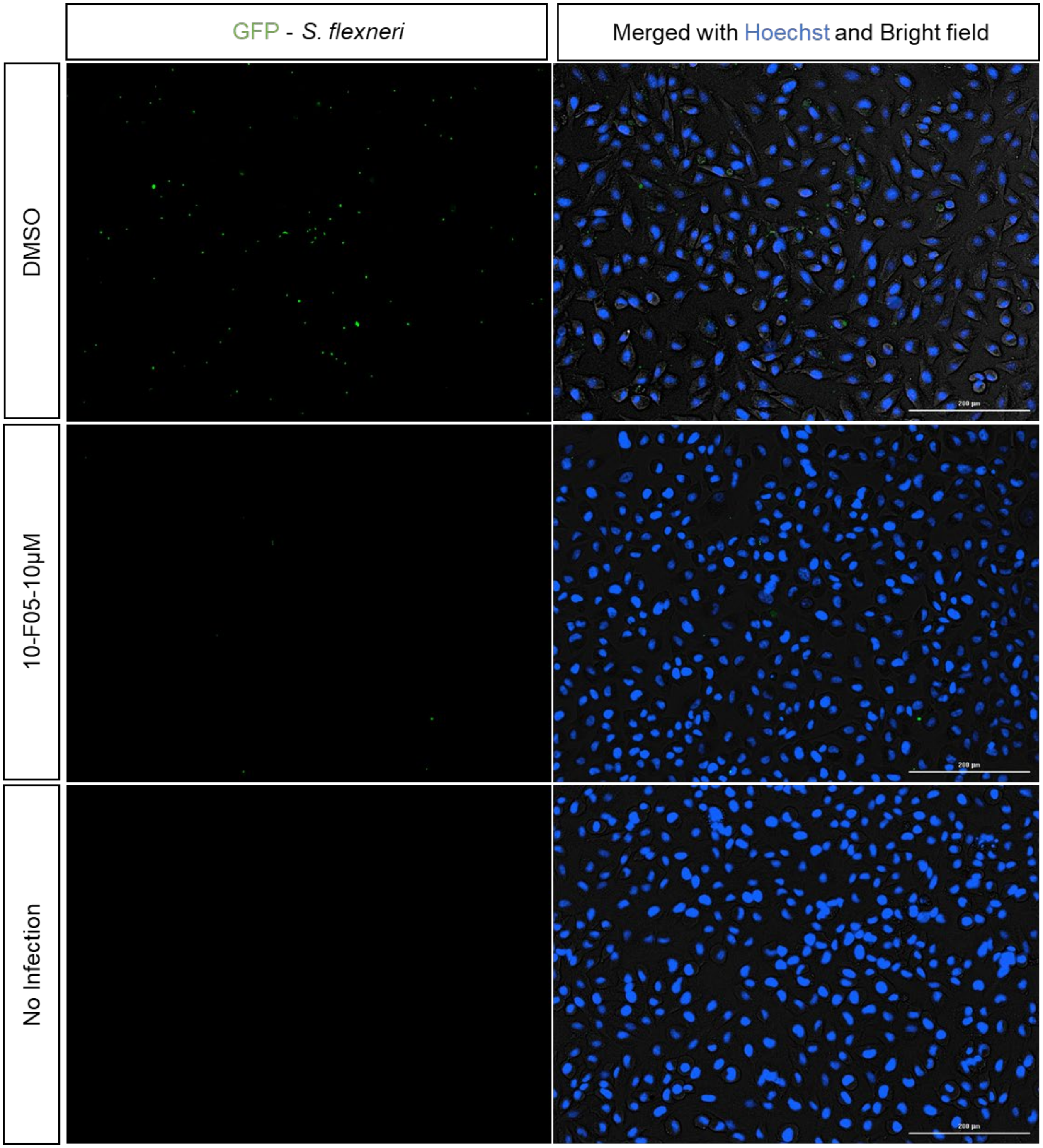

**Figure S14.** Representative images for GFP-labeled *S. flexneri* infection assays in Hela cells. GFP tagged *S. flexneri* M90T was treated with 10 µM of 10-F05 for 1 h before infection. Live cells were stained by Hoechst dye. Scale bar: 200 µm.

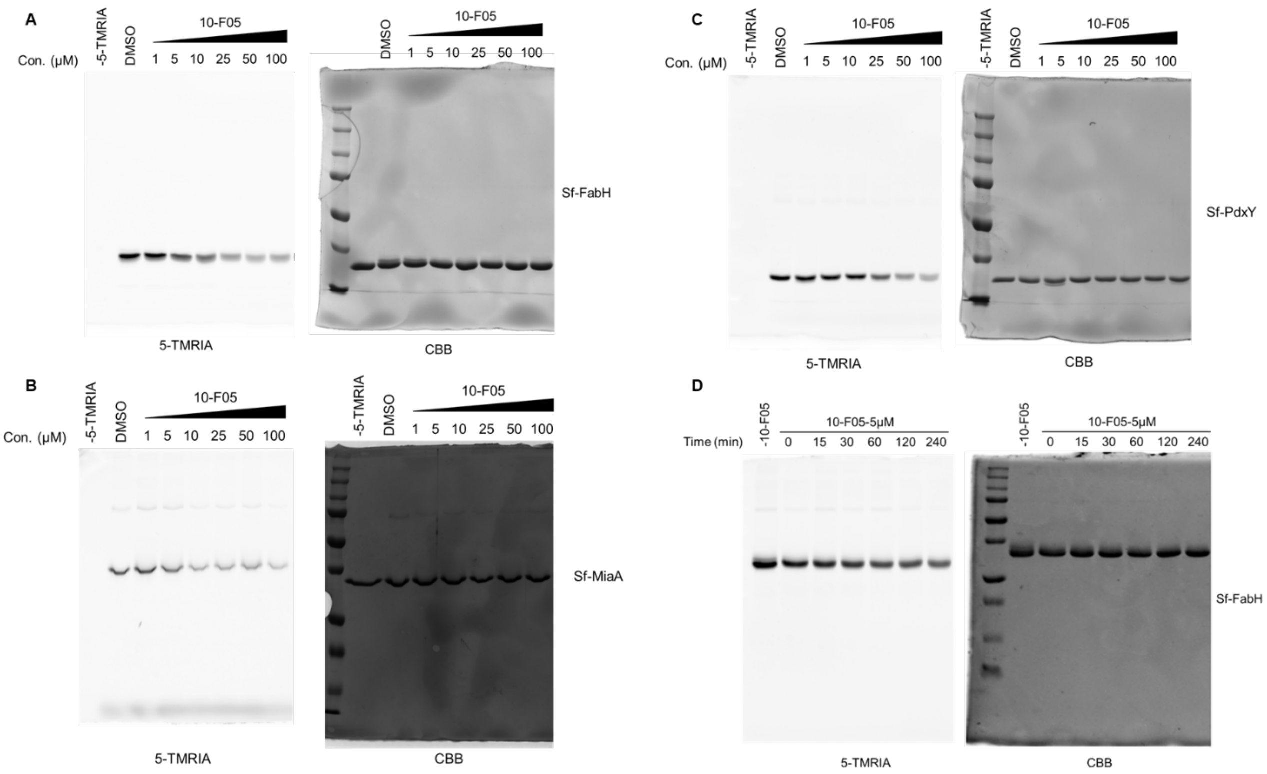

**Uncropped gel for Figure 5A** and **5B.**

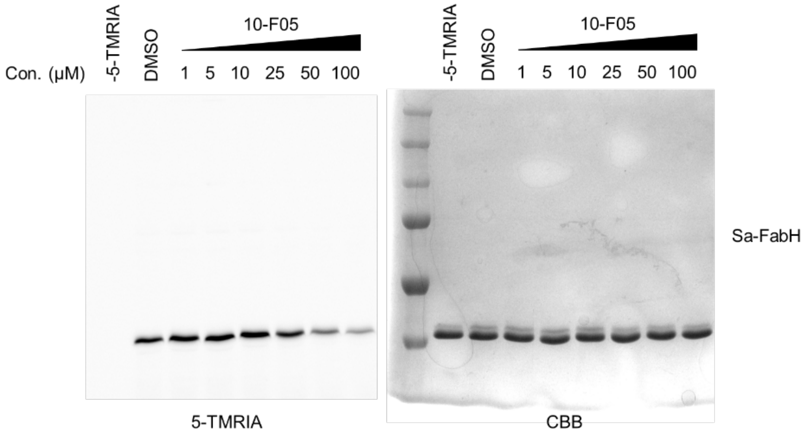

**Uncropped gel for Figure 5C.**

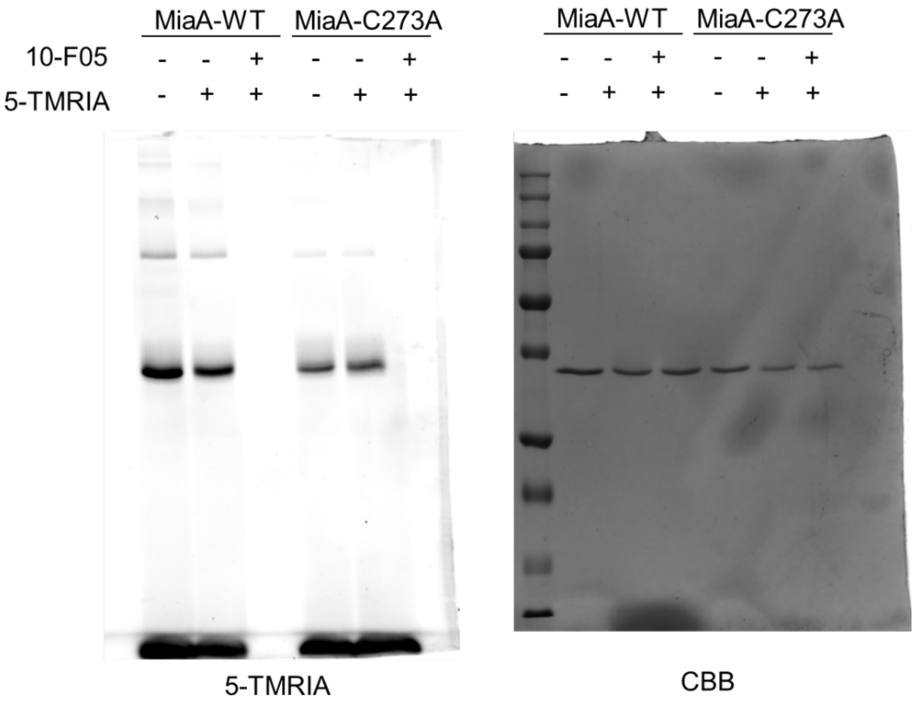

**Uncropped gel for Figure 6A.**

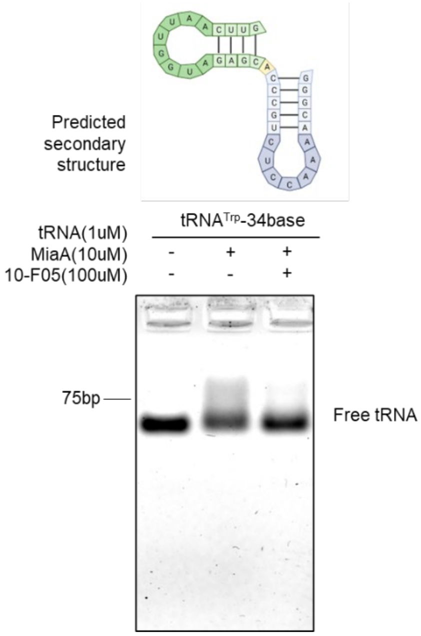

**Uncropped gel for Figure 6C.**

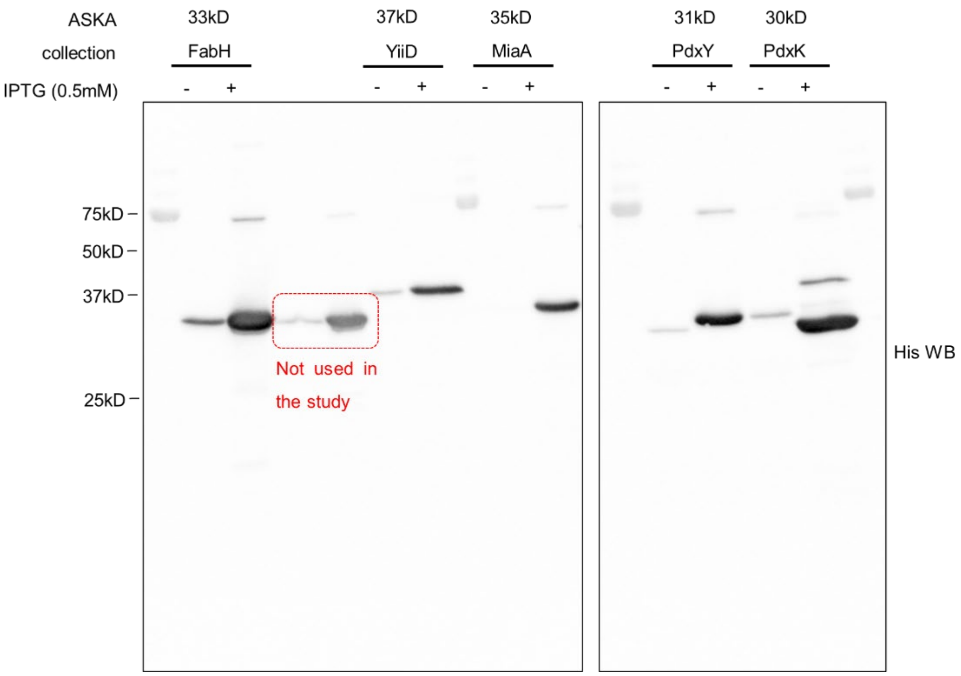

**Uncropped blot for Figure S10.**

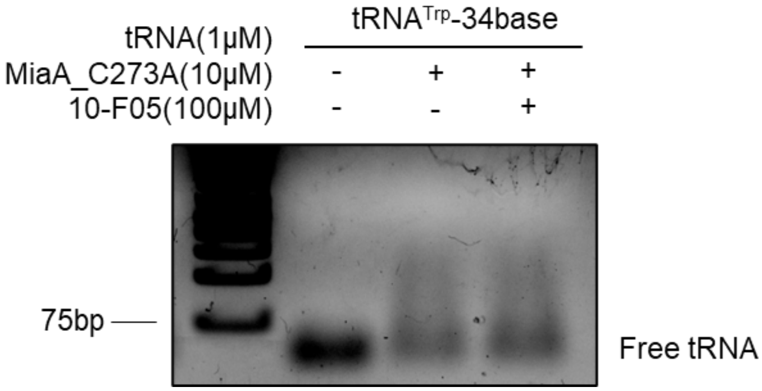

**Uncropped gel for Figure S12.**

### Supplementary Data

Supplementary Data is contained in an Excel file with several tabs. The contents are listed below.

**Tab 1.** Antibacterial Screening Result for the Cys-library.

**Tab 2.** Cytotoxicity Screening Result for the Cys-library.

**Tab 3.** DTNB Reactivity for the Cys-library.

**Tab 4.** Processed ABPP Result

**Tab 5.** Predicted S. flexneri Essential Genes

**Tab 6.** Antibiotic Spectrum

**Tab 7.** Cross Resistance of different antibiotics

**Tab 8.** Cys-library

### Materials and Method

**Reagents, bacterial strains, and cell lines.** Cysteine focused covalent library containing 3200 compounds was purchased from Enamine. Other chemicals used are DMAPP (Dimethylallyl Pyrophosphate, 63180 Cayman Chemical), $N^6$-($\triangle^2$-isopentenyl) adenosine ($i^6A$, 20522, Cayman Chemical), 5,5-dithio-bis-2-nitrobenzoic acid (DTNB, Ellman's Reagent, ThermoFisher), Tetramethylrhodamine-5-Iodoacetamide Dihydroiodide (5-TMRIA, single isomer, ThermoFisher), 2-Methylthio-N-6-isopentenyladenosine ($ms^2i^6A$, sc-484230, Santa Cruz), 10-F05 (MFCD21602491, 1 click chemistry), 10-F05-N (115.316.629, Aurora), 10-L07-N (115.315.060, Aurora), 10-I09-N (ST-0015, Combi-blocks), 10-J03-N(ST-1288, Combi-blocks). All purchased reagents were used as received without further purification unless otherwise noted.

*E. coli* BAA-2340, *A. baumannii* BAA-1790, *P. aeruginosa* BAA-2110, *K. pneumoniae* 700603, *K. pneumoniae* 13883, *S. aureus* MSSA476, *S. aureus* 43300, *E. cloacae* 13047, *E. coli* K12 were purchased directly from ATCC. *V. cholerae* SAD30 was a kind gift from Prof. Tobias Dörr from Cornell University. *S. flexneri* 5a M90T was a kind gift from Prof. Neal M. Alto from UT Southwestern. Enteropathogenic *E. coli* (EPEC) strain JPN15 (serotype O127:H6) was purchased from BEI Resources (NR-50517). *E. coli* strains from Keio collection (JW1077-fabH, JW1628-pdxY, JW3859-yiiD, JW4129-miaA, JW5503-tolC, parent strain BW25113) were purchased from Horizon Discovery. *E. coli* strains from ASKA collection (JW1077-fabH, JW1078-fabD, JW4129-miaA, JW1628-pdxY, JW3859-yiiD, JW5503-tolC, JW2411-pdxK, JW0658-miaB) were from National BioResource Project (https://resourcedb.nbrp.jp/top.jsp).

**MIC measurement.** The minimum inhibitory concentrations (MICs) of compounds were measured using the broth microdilution method.[1] One stab using 10 μL tip from bacteria glycerol stock was inoculated into 5 mL of LB supplemented with selection antibiotics (Kanamycin for Keio collections and chloramphenicol for ASKA collections), and the culture was incubated at 37 °C with shaking overnight. The overnight culture was diluted 1:10000 (~ $5 \times 10^5$ CFU/mL) and then dispensed in to a 96-well plate containing serial dilutions of indicated compounds in DMSO. The plate was incubated at 37 °C for 16 ~ 20 h with shaking at 80 rpm and the $OD_{600}$ was measured using Cytation 5 plate reader (BioTek). MIC was determined as the lowest concentration for which no bacterial growth was observed. LB was used as the growth media for all the bacteria strains unless otherwise mentioned. For *S. aureus* and *A. baumannii* strains, tryptic soy broth (TSB, BD, DF0370-17-3) was used as the growth media instead of LB. For *P. aeruginosa* and *K. pneumoniae* strains, nutrient broth (NB, RPI, N15100) was used as growth media. All MIC measurements were performed in at least biological duplicates.

**Liquid-based antibacterial susceptibility screening.** Bacterial overnight culture was diluted 1:10000 into 50 mL LB (TSB for *S. aureus*). The diluted culture was first dispensed into a 96-well plate (Costar, 96 well flat bottom, 3370) and then transferred (30 µL per well) into 384-well plates (Greiner, 384 well flat bottom, 781101) using epMotion 96 (Eppendorf). Cysteine focused library was diluted into desired stock concentration using DMSO and transferred into the 384-well plates using epMotion96 to a final concentration of 25 µM. The plates were incubated at 37 °C for 16 ~ 20 h with shaking at 80 rpm and the OD600 was measured using Cytation 5 plate reader (BioTek). Bacteria relative growth was calculated by normalizing to the DMSO control group.

**Cloning, protein expression, and purification.** *S. flexneri* FabH, PdxY, MiaA, MiaA_C273A, *S. aureus* FabH in the pET28a (+) vector with an N-terminal His-tag (cloned using the EcoRI and XhoI sites) were purchased from Twist Bioscience. The corresponding plasmid was transformed into BL21(DE3) chemical competent *E. coli* for expression. The transformed *E. coli* was then inoculated from an overnight culture into the 2L of LB supplemented with 50 µg/mL of Kanamycin. The cells were grown at 37 °C for ~4 h with shaking at 200 rpm until the OD600 reached 0.6~0.8. IPTG (0.2 mM) was added to induce protein expression and the culture was incubated at 18 °C overnight with shaking. Cells were harvested by centrifugation (8000 × g, 5 min, 4°C). Bacterial pellets were frozen at −80 °C for future use. Bacterial pellets were resuspended in cold lysis buffer [50 mM Tris pH 8.0, 500 mM NaCl, 0.5 mg/mL Lysozyme (Thermo Scientific, 89833), 1 mM phenylmethylsulfonyl fluoride (PMSF, Thermo Scientific, 36978) in PBS, and Pierce universal nuclease] for 30 min on ice. Then the pellets were sonicated on ice for 2 min at 50% amplitude three times. Lysate was then clarified by centrifugation (30 000 × g, 60 min, 4 °C). The supernatant was first loaded onto pre-equilibrated Ni-NTA resin (Qiagen, 30210), then washed with 50 mL cold wash buffer (50 mM Tris pH 8.0, 500 mM NaCl, 20 mM imidazole), and eluted with cold elution buffer (50 mM Tris pH 8.0, 500 mM NaCl, 200 mM imidazole). The resulting elution was then concentrated using a 10 kD MWCO Amicon filter and fractionated in ÄKTA pure FPLC system using a Superdex 75 gel filtration column pre-equilibrated with protein storage buffer (20 mM Tris, pH 8.0, 60 mM NaCl, 1 mM DTT, 10% glycerol). Fractions containing corresponding proteins were then pooled, flash-frozen, and stored at −80 °C for future use. The protein concentration was measured using a Braford assay (Thermo Scientific, 23200).

**Mammalian cell culture and mammalian cytotoxicity measurement.** Human HEK293T and Hela cells were cultured in DMEM (Invitrogen) media supplemented with 10% (v/v) heat-inactivated fetal bovine serum (FBS, Invitrogen). A549 cells were cultured in RPMI-1640 (Invitrogen) with 10% (v/v) heat-inactivated FBS.

Mammalian cytotoxicity measurement was performed using Cell Titer-Glo 2.0 Cell Viability Assay (Promega, G9243) according to the manufacturer protocol. In brief, mammalian cells were seeded into 384-well white plates (Greiner, 781080) and incubated at 37 °C for one day. Compounds from the Cys-Library were dispensed into the assay plate using epMotion 96. The plates were then incubated at 37 °C for two days. An equal volume of Cell Titer-Glo 2.0 reagent was added into each well of the assay plate using epMotion 96 and incubated at 22 °C for 10 min, followed by luminescence measurement using a Cytation 5 plate reader (BioTek).

**Antibiotic resistance induction.** Resistance induction was performed following a modified version of a previously published method.[2] In brief, the overnight bacterial culture was diluted 1:10000 into fresh LB media (TSB for *S. aureus*) and the MICs of the compounds were detemined. Cultures from the wells with 0.25 × MIC of compounds were diluted 1:100 in fresh media and the MIC measurement was repeated. This procedure was repeated for one month or until the MIC of 10-F05 reached 250 μM. The fold changes in MIC were determined by dividing the daily MIC values by the MIC value on day 1.

**Time-dependent killing assay.** *S. flexneri* M90T overnight culture was diluted 1:10000 into fresh LB media. The starting inoculation (~$10^5$ CFU/mL) was aliquoted into 15 mL tubes containing indicated concentrations of 10-F05. The bacterial culture was subsequently incubated at 37 °C with shaking. Small portions of bacterial culture were collected at indicated time points. Serial dilutions were then plated on agar plates and incubated at 37 °C overnight to determine the CFU. Experiments were performed in biological duplicates.

**Reduced DTNB assay.** Reduced DTNB assay was performed following a previously published method.[3] In brief, a master mix of reduced DTNB was prepared by incubating DTNB (50 μM) with TCEP (200 μM) in DTNB reaction buffer (20 mM sodium phosphate, 150 mM NaCl, pH 7.4) for 5 min at 22 °C. Reduced DTNB was then transferred into a 384-well plate (Greiner, flat bottom, 781101) using epMotion 96 (Eppendorf). Tested compounds (200 μM) were then dispensed into the assay plate, followed by continuous UV measurement at 412 nm at 37 °C in Cytation 5 plate reader (BioTek). The background absorbance at 412 nm was measured under the same conditions without DTNB of each compound and subtracted from corresponding absorbance value. The concentration of remaining $TNB^{2-}$ was calculated based on the absorbance values and normalized to the initial absorbance of each compound. Linear regression using Prism was performed to fit the rate for the first 45 min of measurements.

**In-gel fluorescence analysis.** Purified proteins (1.5 μM) were added into 40 μL of PBS with 10-F05 or DMSO at indicated concentrations. After 1 h incubation at 22 °C or indicated incubation time for time-dependent labeling experiments, 5-TMRIA (10 μM) was added into the solution to label the cysteines. The reaction mixtures were incubated for another 1 h at 22 °C and then quenched by adding 8 μL of 6x Laemmli buffer and analyzed by SDS-PAGE. Proteins labeled by 5-TMRIA were detected using Krypton scanning in ChemiDoc Imaging System (Bio-Rad) and protein loading was measured by Coomassie blue staining. Experiments were performed in biological duplicates and representative images were shown.

**Cell culture and protein labeling for TMTpro18plex-based proteomics.** In brief, overnight culture of indicated bacteria was harvested (4,500 × g, 10 min, 4 °C), washed, and resuspended in 100 μL of PBS to give a final theoretical $OD_{600}$ of 40. Then 50 μM of 10-F05, 10-L07, or DMSO (solvent control) were added into the live bacteria suspension and incubated at 37 °C with shaking for 2 h. Bacterial cells were pelleted (4500 × g, 4 °C, 10 min) and washed with PBS twice. Cells were resuspended in cold solution of DPBS containing Pierce™ Protease and Phosphatase Inhibitor Mini Tablet (ThermoFisher, A32961) or 100 μL Halt™ Protease and Phosphatase Inhibitor Cocktail (ThermoFisher, 78446) (1 tablet or 100 μL per 10 mL), then lysed using a Branson 550 probe sonicator (3 × 10 pulses, 0.4 sec, 40% power, 4 °C). Soluble fractions were then normalized to 2.0 $mg \cdot mL^{-1}$ using the DC Protein Assay (BioRad) and absorbance was measured using a BioTek Cytation 5 plate reader following manufacturer's instructions (BioTek Instruments, Winooski, VT, BTCYT5MPW). Normalized cellular lysates were then treated with desthiobiotin-tagged iodoacetamide probe[4] (100 μM) at ambient temperature for 1 hour by rotating end-over-end (20 rpm). Proteins were precipitated with 600 μL of cold methanol (–20 °C), 200 μL of $CHCl_3$ and 100 μL of chilled water (4 °C). Following centrifugation (15,000 rpm, 10 min, 4 °C), a protein disk formed at the interface of $CHCl_3$ and aqueous layers. Both layers were aspirated without perturbing the disk, which was resuspended in cold methanol (600 μL, –20 °C) and $CHCl_3$ (200 μL, 4 °C) by vortexing and sonicating using a sonicator equipped with a horn cup (1 × Qsonica Q700, Amplitude = 100, Process time = 20 sec, Pulse-ON time = 2 sec, Pulse-OFF time = 1 sec, 4 °C). The proteins were pelleted (15,000 rpm, 10 min, 4 °C), and 100 μL of the digestion buffer (8 M urea, 50 mM TEAB, pH 8.5) was added to the resulting pellets. The pellets were resuspended with sonication and agitated on a thermal mixer (65 °C, 10 min, 1,000 rpm). Then, 5 μL of 200 mM dithiothreitol in water was added to each sample, and the mixture was agitated on a thermal mixer (65 °C, 10 min, 1,000 rpm). Next, 10 μL of 100 mM iodoacetamide in water was added to each sample, and the mixture was agitated on a thermal mixer (37 °C, 30 min, 1,000 rpm).

**Trypsin/Lysine-C digestion and streptavidin enrichment.** 40 μg of Pierce™ Trypsin/Lysine-C Protease Mix (ThermoScientific, MS-Grade A40007) was reconstituted in 60 μL of 50 mM acetic acid and 20 μL of 100 mM calcium chloride. Samples were diluted with 400 μL of 50 mM TEAB (ThermoScientific, 90114) and 4 μL of the Trypsin/Lysine-C solution. Proteins were digested with agitation overnight on a thermal mixer (37 °C, 1,000 rpm). To each sample was then added 500 μL of the enrichment buffer (50 mM TEAB, 0.2% IgepalTM CA-630, pH 8.5) containing 50 μL of Pierce™ Streptavidin agarose resin

(ThermoFisher, 20353). Samples were enriched by rotating end-over-end (20 rpm) for 3 hours at ambient temperature. Samples were next transferred onto Micro Bio-Spin™ Chromatography Columns (Bio-Rad, 7326204) and washed with the wash buffer (3 × 50 mM TEAB, 150 mM NaCl, 0.1% Igepal™ CA-630), DPBS (3×), and water (3×) by carefully aspirating from the bottom of each Bio-Spin column without drying the resin. Peptides were eluted with 50% acetonitrile in water containing 0.1% formic acid and each sample evaporated to dryness using vacuum centrifugation overnight (Savant, SpeedVac SPD-2030, temperature = 40 °C, vacuum pressure = 5.1 Torr).

**TMTpro-18plex labeling.** Peptides were redissolved in 100 µL EPPS buffer (200 mM, pH 8.5) with 30% acetonitrile. TMT tags (10 µL per channel in acetonitrile, 20 $\mu g \cdot \mu L^{-1}$) were added to the corresponding tubes and agitated on a thermal mixer (25 °C, 90 min, 1,000 rpm). Each reaction was quenched by the addition of 10 µL of 5% hydroxylamine and mixed (25 °C, 15 min, 1,000 rpm). To each sample, 10 µL of formic acid was added and mixed (25 °C, 5 min, 1,000 rpm). TMT-labeled samples were combined into a single Protein LoBind microcentrifuge tube and evaporated to dryness using vacuum centrifugation.

**Peptide desalting.** Sep-Pak® C18 cartridges (Waters, WAT054955) were conditioned with acetonitrile (3×) and desalting buffer (3×, 95% water, 5% acetonitrile, 0.5% formic acid). TMT-labeled peptides were redissolved in 500 µL of the desalting buffer, loaded dropwise onto the cartridge, and eluted at the rate of approximately 1 drop per second. The cartridge was then reloaded with the flow-through and subsequently desalted by slowly passing desalting buffer (3 × 1 mL). The peptides were eluted by adding 500 µL of 80% acetonitrile, 20% water, 0.5% formic acid (3×), eluates were combined into a clean Protein LoBind microcentrifuge tube, and sample was evaporated to dryness using vacuum centrifugation.

**High pH reverse-phase fractionation.** The spin columns for high pH fractionation (Pierce high pH reverse-phase peptide fractionation kit, ThermoScientific, 84868) were pre-equilibrated according to manufacturer's instructions prior to use. Desalted peptides were redissolved in 0.1% trifluoracetic acid aqueous solution and loaded onto the column. The columns were spun down (2,000 × g, 2 min) with eluate retained, washed with 300 µL of water with eluate retained, and subjected to fractionation with a series of 0.1% triethylamine/acetonitrile buffers (2,000 × g, 2 min) with each eluate collected into a clean Protein LoBind tube. The following buffers were used for peptide elution (% acetonitrile): 5, 7, 9, 11, 12, 13, 14, 15, 16, 17, 18, 19, 20, 21, 22, 23, 24, 25, 26, 27, 28, 29, 30, 35, 40, 45, 50, 80. Fractions were evaporated to dryness using vacuum centrifugation, resuspended in water, and peptide concentrations determined using NanoDrop One Spectrophotometer (ThermoScientific, ND-ONEC-W, version 2.2.0.16). Fractions were combined with at least 7 fractions separation to yield 10 total fractions with approximately equivalent peptide amounts, filtered through CoStar Spin-X columns (Corning, 8160) and evaporated to dryness. The resulting 10 fractions were each reconstituted in 62 µL of 2% acetonitrile with 0.5% formic acid for subsequent nanoLC-MS/MS analysis.

**Nano-scale reverse phase chromatography and tandem MS (nanoLC-MS/MS)**. The nanoLC-MS/MS analysis was carried out using an Orbitrap Eclipse (ThermoScientific, San Jose, CA) mass spectrometer equipped with a nanospray Flex Ion Source coupled with the UltiMate 3000 RSLCnano (Dionex, Sunnyvale, CA). Each reconstituted fraction (3.5 mL = 0.7 µg for global proteomics fractions) was injected onto a PepMap C-18 RP nano trap column (5 µm, 100 µm × 20 mm, Dionex) at 20 $\mu L \cdot min^{-1}$ flow rate for rapid sample loading, and separated on a PepMap C-18 RP nano column (2 µm, 75 µm × 25 cm). The column was equilibrated with 2% acetonitrile in 0.1% aqueous formic acid (eluant A) prior to each run. The labeled peptides were eluted in a 120-min gradient of 5% to 33% eluant B containing 95% acetonitrile in 0.1% formic acid at 300 $nL \cdot min^{-1}$, followed by an 8-min ramping to 90% B, a 7-min hold and 21-min re-equilibration with 2% acetonitrile and 0.1% formic acid prior to the next run. The Orbitrap Eclipse was operated in positive ion mode with nano spray voltage set at 1.9 kV and source temperature at 300 °C. External calibration for FT, IT and quadrupole mass analyzers were performed. Raw MS data files for all the fractions were acquired using a real-time search (RTS) synchronous precursor selection (SPS) $MS^3$ workflow as reported previously.[5] Specifically, the RTS MS3 workflow consisted of 2.5 second "Top Speed" data-dependent CID-MS/MS scans (for peptide identifications by RTS) that enabled to trigger SPS of 10 MS2 product ions for subsequent MS3 in FT. In RTS node, the *Homo sapiens* FASTA database containing 20,520 sequences was imported along with trypsin as the enzyme for real-time spectral database search for the samples from corresponding species. The search parameters included: TMTpro modification on N-terminal amines (D mass 304.2071) and carbamidomethyl modification of cysteine (D mass 57.0215) as static modifications; TMTpro modification (D mass 304.2071) on lysine, desthiobiotin-tagged STP ester probe (D mass 196.1212), and methionine oxidation (D mass 15.9949) as dynamic modifications; maximum 3 variables per peptide; and 2 maximum missed cleavage allowed. A maximum search time of 35 ms was allowed for the RTS MS3 searching. The MS3 scan was carried out using a mass range of 110-500 m/z, an MS isolation window of 1.1 m/z and MS2 isolation window of 2.0 m/z were used. A resolving power of 50,000 at MS3 with a normalized collision energy of 55% was used for peptide quantitation. Other parameters included 200% normalized AGT target and 120 ms for maximum injection time. Dynamic exclusion parameters were set at 1 count within 50s exclusion duration with ±10 ppm exclusion mass window. All data were acquired under Xcalibur 4.4 operation software in Orbitrap Eclipse (ThermoScientific, San Jose, CA).

**Data processing, protein identification, and data analysis**. All raw MS spectra were processed and searched using the Sequest HT search engine within the Proteome Discoverer 3.0 (PD3.0,

ThermoScientific). The same database for human proteins used for RTS data acquisition as described above was used for post-MS database searches. The default search settings used for 18-plex TMT quantitative processing and protein identification in PD3.0 searching software were: two mis-cleavage for full trypsin with fixed carbamidomethyl modification of cysteine, fixed 18-plex TMT modifications on lysine and N-terminal amines along with variable modifications of methionine oxidation, and protein N-terminal acetylation. The peptide mass tolerance and fragment mass tolerance values were 10 ppm for MS survey scan, 0.6 Da for MS2 and 20 ppm for MS3, respectively. Identified peptides were further filtered for maximum 1% FDR using the Percolator algorithm in PD3.0 along with additional peptide confidence set to high and peptide mass accuracy ≤5 ppm. The TMT18-plex quantification method within Proteome Discoverer 3.0 software was used to calculate the reporter ion abundances in MS3 spectra that were corrected for the isotopic impurities. Both unique and razor peptides were used for relative protein quantitation. Signal-to-noise (S/N) values were used to represent the reporter ion abundance with a co-isolation threshold of 50% and an average reporter S/N (intensity) threshold of ≥10 used for quantitation spectra. The intensities of peptides, which were summed from the intensities of the PSMs, were summed to represent the abundance of the proteins. For relative ratio between the two groups, normalization on sum of total peptide intensities for each sample was applied. The search results including ratio, peptide abundance for each sample were output to Microsoft Excel software for further data analysis.

**tRNA preparation.** 34-nucleotide RNA oligomers were purchased from Integrated DNA Technology (idtdna.com). The sequence used in the study is: $tRNA^{Trp}$-34bases, [GUUCAAUUGGUAGAGCACCGGUCU-CCAAAACCGG]. RNA oligomers were reconstituted in tRNA folding buffer (30 mM HEPES, 100 mM KCl, 2 mM $MgCl_2$, and 50 mM ammonium acetate pH 7.0). To ensure homogeneous folding of RNA into predicted structure with two hairpin loops, 50 µL of RNA solutions were heated up to 85 °C over the course of 2 min and then cooled to 4 °C over the course of 35 min in a PCR thermal cycler (Applied Biosystems Veriti 96-well Thermal Cycler). tRNA secondary structure prediction was performed as previously described.[6]

**MiaA binding assay.** MiaA binding assay was performed following a previously published method with modifications.[7] Purified MiaA (10 µM) was incubated with 10-F05 (100 µM) or DMSO in 40 µL tRNA binding buffer (50 mM Tris-HCl pH 8.0, 100 mM NaCl, 10 mM $MgCl_2$) for 1 h at 22 °C. The folded tRNA (1 µM) was added into the mixture and incubated for another 20 mins. The reaction mixtures were then supplemented with 8 µL of 6x Tri tracker loading dye (ThermoFisher) and separated on a 2 % agarose TAE gel pre-stained with GelRed (Biotium, 41003) for 20 min at 100 V. Gels were then visualized in ChemiDoc Imaging System (Bio-Rad) and analyzed using ImageJ (1.53t). The unbound tRNA ratio was calculated by normalizing to the intensity of tRNA-only group. Experiments were performed in biological quadruplicates.

**MiaA activity assay.** MiaA activity assay was performed following a previously published method[8]. Purified MiaA (200 nM) was incubated with 10-F05 (50 μM) or DMSO in 800 μL TMD buffer (60 mM Tris-HCl pH 7.5, 20 mM $MgCl_2$, and 2 mM DTT) at 22 °C for 1 hour. Dimethyl allyl pyrophosphate (100 μM, DMAPP), bovine serum albumin (100 μg, BSA) and folded tRNA (6 μM) were then added to isopentenylate the A37 residue. Sample aliquots (60 μL) were removed at selected time points (0−30 min), heat-denatured at 95 °C for 5 min. Nuclease P1 (M0660S, NEB, 200 units/mL in 30 mM sodium acetate buffer pH 5.4) and $ZnSO_4$ (10 mM) were immediately added into the cooled aliquots and incubated with shaking for 16 h at 37 °C. 10X CutSmart Buffer (NEB, B6004) and quick CIP (NEB, M0525S, 1 μL of 5000 units/mL) were added into the aliquots and incubated for 4 h at 37 °C to yield the ribonucleosides. Proteins were removed by adding 50 μL of cold acetonitrile and centrifugation (17000 × g, 10 min, 22 °C). Supernatants were collected for LC-MS analysis. The reaction components were eluted at a flow rate of 0.3 mL/min with the following time program: 0 % B from 0 to 2 min, 0 % B to 100 % B from 2 to 5 min, 100 % B from 5 to 10 min, 100 % B to 0 % B from 10 min to 12 min. Buffer A was 0.1 % formic acid in HPLC-water and buffer B was 0.1 % formic acid in HPLC-acetonitrile. The mass spectrometer was operated in positive ion mode. The observed m/z value of the $[M+H]^+$ stage of synthetic $i^6A$ standard (Cayman) was 336.2833 and the retention time was 5.27 min. Serial dilutions (0.01 ~ 10 μM) of synthetic standard $i^6A$ were prepared in 50 % acetonitrile and 50% water and injected into LC-MS to generate the standard curve. Concentrations of $i^6A$ in each sample were then calculated based on the standard curve and the initial velocities were calculated using linear regression analysis in GraphPad Prism9. Experiments were performed in biological duplicates.

**Molecular modeling.** A reference MiaA structure from *E. coli* was downloaded from PDB database (2ZM5) and loaded into MOE (2020) software as Biomolecule Assembly with default settings. The protein structure was then prepared using the QuickPrep function with the default parameters and thoroughly checked using the Structure Preparation function. The tRNA was first removed from the binding pocket. The binding site on MiaA was defined based on the identified cysteine position. 10-F05 was docked onto Cys273 using Covalent Docking (Reaction: alpha-halocarbonyl, thioether; Refinement: Induced fit). The resulting docking pose was then superposed with the tRNA-bound MiaA structure and visualized in MOE and PyMol (4.6.0).

**Frame shift quantification.** Frame shift quantification was performed as previously described.[9] The plasmids we used are kind gifts from Dr. Matthew A. Mulvey. Briefly, *E. coli* WT and *miaA* KO strains (Keio collection) transformed with pCWR44 and pCWR45 were grown overnight in LB supplemented with chloramphenicol for WT strains and both chloramphenicol (25 μg/mL) and kanamycin (50 μg/mL) for *miaA* KO strains. The cells were sub-cultured into fresh LB to give an $OD_{600}$ of 0.1 at the day of experiment. 5 μM of 10-F05 was added into the bacterial culture and incubated at 37 °C with shaking. After 30 min, arabinose (0.2%) was added into the bacterial cultures to induce the expression of luciferase. Bacterial cells were then pelleted when the $OD_{600}$ reached 0.5 and resuspended in Passive Lysis Buffer (Promega, E1910), followed by mechanical lysis using disruption beads in TissueLyser LT (QIAGEN). Luciferase activities were analyzed using the dual luciferase reporter assay following the manufacturer protocol (Promega, E1910). Experiments were performed in biological triplicates. Translation error was calculated based on the following formula:

$$Translation\ error(\%) = \frac{LumFirefly(pCWR44)}{LumRenilla(pCWR44)} \div \frac{LumFirefly(pCWR45)}{LumRenilla(pCWR45)} \times 100(\%)$$

**Quantification of $ms^2i^6A$-tRNA modification.** tRNA purification was performed following a previously published method.[10] In brief, bacterial overnight culture (WT and *miaA* KO *E. coli* strains) was diluted 1:10 into fresh media (25 mL) containing 100 μM of 10-F05 or DMSO as control. After 4 h incubation at 37 °C with shaking, the bacterial cells were pelleted by centrifugation (4500 × g, 25 min, 4°C). The pellets were washed with 0.9% NaCl once and stored at -80 °C for future use. Total nucleic acids were extracted from the pellet using 900 μL of extraction buffer (50 mM NaOAc, 10 mM MgOAc, pH 5.0) and 860 μL of acidic phenol pH 4.5 (Sigma, P4682) for 30 min at 37 °C with shaking. The aqueous phases were collected after centrifugation (4500 × g, 15 min, 4 °C). Next, 700 μL of extraction buffer was added into the phenol phase and the same procedures were repeated. The aqueous phases were combined. 75 μL of 5 M NaCl and 1.5 mL of isopropanol was added into the combined aqueous phases to precipitate the total nucleic acids by centrifugation (14500 × g, 15 min, 22 °C). The pellets were washed with cold 70 % ethanol and air dried for 10 min. rRNA was removed by resuspending the pellets into 750 μL of cold 1 M NaCl and centrifugation (9500 × g, 20 min, 4 °C). Remaining nucleic acids (DNA and tRNA) were precipitated by adding 1.5 mL of cold ethanol to the supernatants and incubated at –20 °C for 30 min, followed by centrifugation (14500 × g, 5 min, 4 °C). The pellets were washed with 70 % cold ethanol and air dried for 10 min. DNA was removed by dissolving the pellets into 300 μL of 0.3 M NaOAc, pH 5.0 and precipitated by adding 170 μL of isopropanol, followed by incubation at 22 °C for 10 min. The supernatants containing tRNA were collected after centrifugation (14500 × g, 5 min, 22 °C). Next, 115 μL of isopropanol was added into the supernatant and incubated at –20 °C for 30 min. The total tRNA was then prepared by centrifugation (14500 × g, 15 min, 4 °C) and washed with 70 % ethanol. A total of 25 μL DEPC-water was added to dissolve the tRNA pellet for each sample. The concentration and quality of tRNA samples were determined by Nanodrop.

Quantification of $ms^2i^6A$-tRNA modification was performed using LC-MS following a previously published method.[11] In brief, approximately 12.5 mg tRNA samples were enzymatically digested into nucleosides in a total of 50 μL of digestion buffer [1 μL of Pierce Universal Nuclease (ThermoFisher, 88700), 0.5 μL of Nuclease P1 (NEB, M0600S, 5000 units/mL), 6 μL of Nuclease P1 buffer and 1 μL of Alkaline phosphatase (Sigma, P5521, 1000 units/mL)] and incubated at 37 °C with shaking overnight. The reaction components were eluted at a rate of 0.3 mL/min with the following time program: 0 % B from 0 to 2 min, 0 % B to 100 % B from 2 to 5 min, 100 % B from 5 to 10 min, 100 % B to 0 % B from 10 min to 12 min. Buffer A is 0.1 % formic acid in HPLC-water and buffer B is 0.1 % formic acid in HPLC-acetonitrile. The mass spectrometer was operated in positive ion mode. The observed m/z value of the $[M+H]^+$ stage of synthetic $ms^2i^6A$ standard (Santa Cruz) was 382.2901 and the retention time was 5.65 min. The normalized $ms^2i^6A$ peak intensity was calculated by dividing the absolute intensity of the $ms^2i^6A$ ions to the tRNA concentration of each sample measured by Nanodrop. Experiments were performed in biological triplicates.

**Osmotic stress resistance assays.** Bacterial overnight culture was diluted 1:1000 into 3% NaCl-LB or LB as indicated. 200 μL of diluted cultures were then dispensed into 96 well plates with 10 μM of 10-F05 or DMSO (control). $OD_{600}$ curve was monitored using a Cytation5 plate reader every 10 mins at 37 °C with consistent shaking. Experiments were performed in biological triplicates.

**Infection of mammalian cells with GFP-tagged S. flexneri M90T.** GFP-tagged *S. flexneri* infection was performed as previously described with modifications.[12] In brief, ~$10^5$ Hela cells were seeded into 24-well plates. An overnight culture of GFP-tagged *S. flexneri* M90T was diluted 100 fold in fresh LB and incubated at 37 °C with shaking until the $OD_{600}$ reached 0.4. The bacterial cells were then pelleted and resuspended in the same volume of PBS. The bacterial suspensions were treated with 10 μM of 10-F05 or DMSO for 1 h at 37 °C with shaking. A small portion of the mixture was plated on LB agar plates overnight to count the CFU after drug treatment. 25 μL of bacterial suspensions were added into Hela cells for an MOI of ~25. After 2 h incubation at 37 °C, the Hela cells were washed with PBS twice and incubated in fresh media containing 50 μg/ml gentamycin and Hoechst statin (Invitrogen, 33342) for 30 min. Hela cells were then washed with PBS twice and imaged using Cytation5. For the agar plates counting assay, Hela cells were directly collected after 30 min incubation and lysed using 0.5% Triton X-100 PBS lysis buffer to release the intracellular bacteria. Serial dilutions from the lysates were plated on agar plates using spot format and incubated at 37 °C overnight. Experiments were performed in biological duplicates.

**Structural based sequence alignment.** Protein structures were downloaded from Uniprot Database using either reported structures or AlphaFold2 predicted structures. Structure based sequence alignment was performed using PROMALS3D (http://prodata.swmed.edu/promals3d/promals3d.php).[13]

**References.**

1 Zanon, P. R. A., Lewald, L. & Hacker, S. M. Isotopically Labeled Desthiobiotin Azide (isoDTB) Tags Enable Global Profiling of the Bacterial Cysteinome. *Angew Chem Int Ed Engl* **59**, 2829-2836, doi:10.1002/anie.201912075 (2020).

2 Bakker, A. T. *et al.* Chemical Proteomics Reveals Antibiotic Targets of Oxadiazolones in MRSA. *J Am Chem Soc*, doi:10.1021/jacs.2c10819 (2022).

3 Resnick, E. *et al.* Rapid Covalent-Probe Discovery by Electrophile-Fragment Screening. *J Am Chem Soc* **141**, 8951-8968, doi:10.1021/jacs.9b02822 (2019).

4 Kuljanin, M. *et al.* Reimagining high-throughput profiling of reactive cysteines for cell-based screening of large electrophile libraries. *Nat Biotechnol* **39**, 630-641, doi:10.1038/s41587-020-00778-3 (2021).

5 Fu, Q. *et al.* Comparison of MS(2), synchronous precursor selection MS(3), and real-time search MS(3) methodologies for lung proteomes of hydrogen sulfide treated swine. *Anal Bioanal Chem* **413**, 419-429, doi:10.1007/s00216-020-03009-5 (2021).

6 Lorenz, R. *et al.* ViennaRNA Package 2.0. *Algorithms for molecular biology : AMB* **6**, 26, doi:10.1186/1748-7188-6-26 (2011).

7 Nozawa, K. *et al.* Crystal structure of Cex1p reveals the mechanism of tRNA trafficking between nucleus and cytoplasm. *Nucleic Acids Res* **41**, 3901-3914, doi:10.1093/nar/gkt010 (2013).

8 Subedi, B. P., Corder, A. L., Zhang, S., Foss, F. W., Jr. & Pierce, B. S. Steady-state kinetics and spectroscopic characterization of enzyme-tRNA interactions for the non-heme diiron tRNA-monooxygenase, MiaE. *Biochemistry* **54**, 363-376, doi:10.1021/bi5012207 (2015).

9 Fleming, B. A. *et al.* A tRNA modifying enzyme as a tunable regulatory nexus for bacterial stress responses and virulence. *Nucleic Acids Res* **50**, 7570-7590, doi:10.1093/nar/gkac116 (2022).

10 Avcilar-Kucukgoze, I., Gamper, H., Hou, Y. M. & Kashina, A. Purification and Use of tRNA for Enzymatic Post-translational Addition of Amino Acids to Proteins. *STAR Protoc* **1**, 100207, doi:10.1016/j.xpro.2020.100207 (2020).

11 Su, D. *et al.* Quantitative analysis of ribonucleoside modifications in tRNA by HPLC-coupled mass spectrometry. *Nat Protoc* **9**, 828-841, doi:10.1038/nprot.2014.047 (2014).

12 Wang, M. *et al.* Golgi stress induces SIRT2 to counteract Shigella infection via defatty-acylation. *Nat Commun* **13**, 4494, doi:10.1038/s41467-022-32227-x (2022).

13 Pei, J., Kim, B. H. & Grishin, N. V. PROMALS3D: a tool for multiple protein sequence and structure alignments. *Nucleic Acids Res* **36**, 2295-2300, doi:10.1093/nar/gkn072 (2008).
